# Dopamine responses reveal efficient coding of cognitive variables

**DOI:** 10.1101/2020.05.20.100065

**Authors:** Asma Motiwala, Sofia Soares, Bassam V. Atallah, Joseph J. Paton, Christian K. Machens

## Abstract

Reward expectations based on internal knowledge of the external environment are a core component of adaptive behavior. However, internal knowledge may be inaccurate or incomplete due to errors in sensory measurements. Some features of the environment may also be encoded inaccurately to minimise representational costs associated with their processing. We investigate how reward expectations are affected by differences in internal representations by studying rodents’ behaviour and dopaminergic activity while they make time based decisions. Several possible representations allow a reinforcement learning agent to model animals’ choices during the task. However, only a small subset of highly compressed representations simultaneously reproduce, both, animals’ behaviour and dopaminergic activity. Strikingly, these representations predict an unusual distribution of response times that closely matches animals’ behaviour. These results can inform how constraints of representational efficiency may be expressed in encoding representations of dynamic cognitive variables used for reward based computations.

## Introduction

The theory of reinforcement learning (RL) provides a large and growing set of algorithms by which animals may learn to interact with their environment using reward feedback. A key component in many RL algorithms is a reward prediction error (RPE) signal that drives learning via the algorithm of temporal-difference (TD) learning (Dayan & Sejnowski, 1994; Sutton, 1988). Correlates of such TD RPEs have been found in the phasic activity of dopaminergic neurons in the midbrain (Bayer & Glimcher, 2005; Fiorillo et al., 2003; Schultz et al., 1997), and electrical and optogenetic manipulations of midbrain dopamine neurons have demonstrated that dopamine neuron activity can function to teach animals about the value of actions (Reynolds et al., 2001; Stauffer et al., 2014; Steinberg et al., 2013). These data have provided compelling evidence that neural systems function similarly to TD RL algorithms. Indeed, a large body of research on dopaminergic signalling supports the hypothesis that reward-based decision-making implemented in neural circuits is well described by the framework of RL (see Niv & Langdon, 2016; Watabe-Uchida et al., 2017 for reviews).

A key problem in explaining dopaminergic activity (DA) in terms of RPEs is that RPEs depend on animals’ expectations that are computed from internal representations of any given environment. However, often, the nature of internal representations used to guide reward based behaviour can not be directly characterised. Understanding how animals construct internal representations of the environment to guide adaptive behavior is, in general, a key outstanding goal of cognitive and systems neuroscience. Within the RL framework, the nature of internal representations places constraints on both reward expectations and RPE signals that may be encoded in neural circuits (Daw et al., 2006; Ludvig et al., 2008; Suri & Schultz, 1998). Thus, examining activity of midbrain DA neurons in terms of RPEs during carefully controlled tasks can be a powerful means for discovering the principles that describe internal representations used by animals to guide cognition and behavior (Botvinick, 2008; Gershman, Norman, & Niv, 2015; Russek, Momennejad, Botvinick, Gershman, & Daw, 2017).

As an example of such a task, we studied animals making decisions based on internal estimates of elapsed time. Previous work has shown that population activity in cortical and striatal circuits in time-based tasks show rich sequential activity (Gouvêa et al., 2015; Mello et al., 2015; Remington et al., 2018; J. Wang et al., 2018). We constructed RL agents that encode internal representations consistent with these observed patterns of activity and trained them on an interval discrimination task on which animals were also trained. We examine how changing different aspects of the representations used can yield different predictions about RPEs and their relation to behavior on a trial-by-trial basis. We find that only agents with internal representations that were inaccurate in encoding the task along a particular dimension showed RPEs that match the profile of dopaminergic activity recorded in mice, and its relation to behavior. Moreover, TD learning using this compressed representation predicted procrastination of choices for a subset of stimulus estimates that closely matched animals’ behaviour.

Representational efficiency has been extensively studied in sensory systems (Atick & Redlich, 1992; Lewicki, 2002; Olshausen & Field, 1996; Rieke, Bodnar, & Bialek, 1995), and has also been shown to be subject to behavioural salience (Machens et al., 2005; Reinagel & Zador, 1999; Salinas, 2006). Based on these results, it has been proposed that constraints of representational efficiency are very likely to affect animals’ reward expectations as well (Botvinick, Weinstein, Solway, & Barto, 2015). However, what redundancies in variables needed for reward based computations are being exploited to achieve efficient representations and how such representations affect reward expectations is still an open question. Our results provide empirical support for the strategy where representational efficiency is achieved such that only the overall number of rewards obtained is preserved (or temporal-difference reward prediction errors are minimised), a finding with wide reaching implications for the more general problem of understanding the neural mechanisms underlying cognition.

## Results

We analysed behaviour and dopaminergic activity of mice performing a time interval discrimination task (Figure 1a). On each trial, animals indicated whether the interval between two tones was longer or shorter than 1.5 seconds. Animals reported their decisions for long’ or short’ intervals in one of two choice ports. For the longest and shortest intervals, animals almost always chose the correct port, but as intervals approached the decision boundary, choices became more variable as captured by animals’ psychometric functions (Figure 1b).

**Figure 1:**
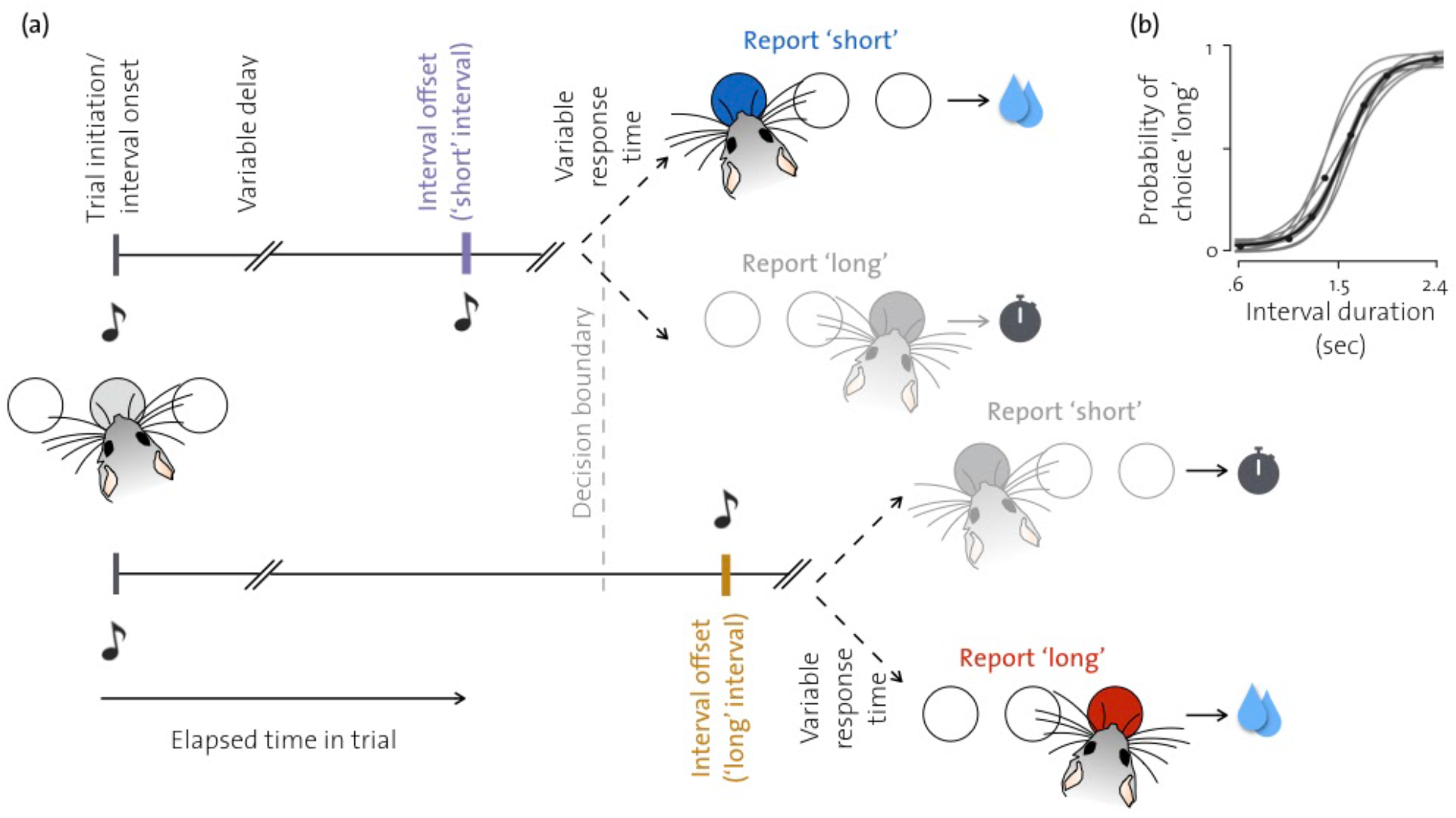
Rodents were trained to classify interval durations. **(a)** Schematic illustrating the timeline of the main events during the interval discrimination task. Animals are presented with three ports and are required to initiate each trial in the central port. The central nose poke triggers a tone and after a variable interval a second tone is presented. These intervals can either be longer or shorter than the decision boundary. Animals have to report ‘short’ or ‘long’ judgements in the two lateral ports based on their estimate of the time elapsed between the two tones. Correct choices result in water reward and incorrect choices result in a time out. **(b)** Psychometric curves of animals performing the task. Grey lines indicate sigmoid fits to behaviour of individual animals (n=6), and black line and dots indicate average over all animals. (Panel (b) adapted from Soares et. al. 2016)

Previous work has shown that animals’ estimates of elapsed time vary from trial to trial. Moreover, neural correlates of such variability have been found in the population activity of striatal circuits (Gouvêa et al., 2015), and the activity of midbrain dopaminergic neurons in the SNc (Soares et al., 2016). However, the structure of internal representations that would be required to explain dopaminergic RPEs in relation to behavior during this task is unknown. To address this, we built a number of RL models that vary in the internal representations of the task environment they use to compute reward expectations. We compared both behavior and RPE’s produced by those models to the behavior of mice and activity from genetically identified dopaminergic neurons in the substantia nigra pars compacta (SNc) using fiber photometry (see Soares et al., 2016).

### Reward expectations are modulated by internal and external states

Reward prediction errors of the form used in TD-learning can arise due to discrepancies between the probability, amount, or timing of expected and actual rewards, or if an unpredictable change in the environment leads to a change in expected future rewards, e.g., by observing an unpredictable reward-predicting cue. During the interval discrimination task the timing of trial onset was unpredictable (Soares et al., 2016). Hence the cue at interval/trial onset, which predicts a potentially forthcoming reward, should thereby cause changes in animals’ reward expectations and concomitant RPEs. Consistent with this reasoning, we found that the tone marking interval/trial onset elicits a phasic DA response (Figure 2a). Since the tone at interval onset was identical for all trials, the phasic DA response was not modulated by stimulus identity. We see that the first peak in the average DA response, after the grey tick on the x-axis of figure 2a, is not modulated by interval duration.

**Figure 2:**
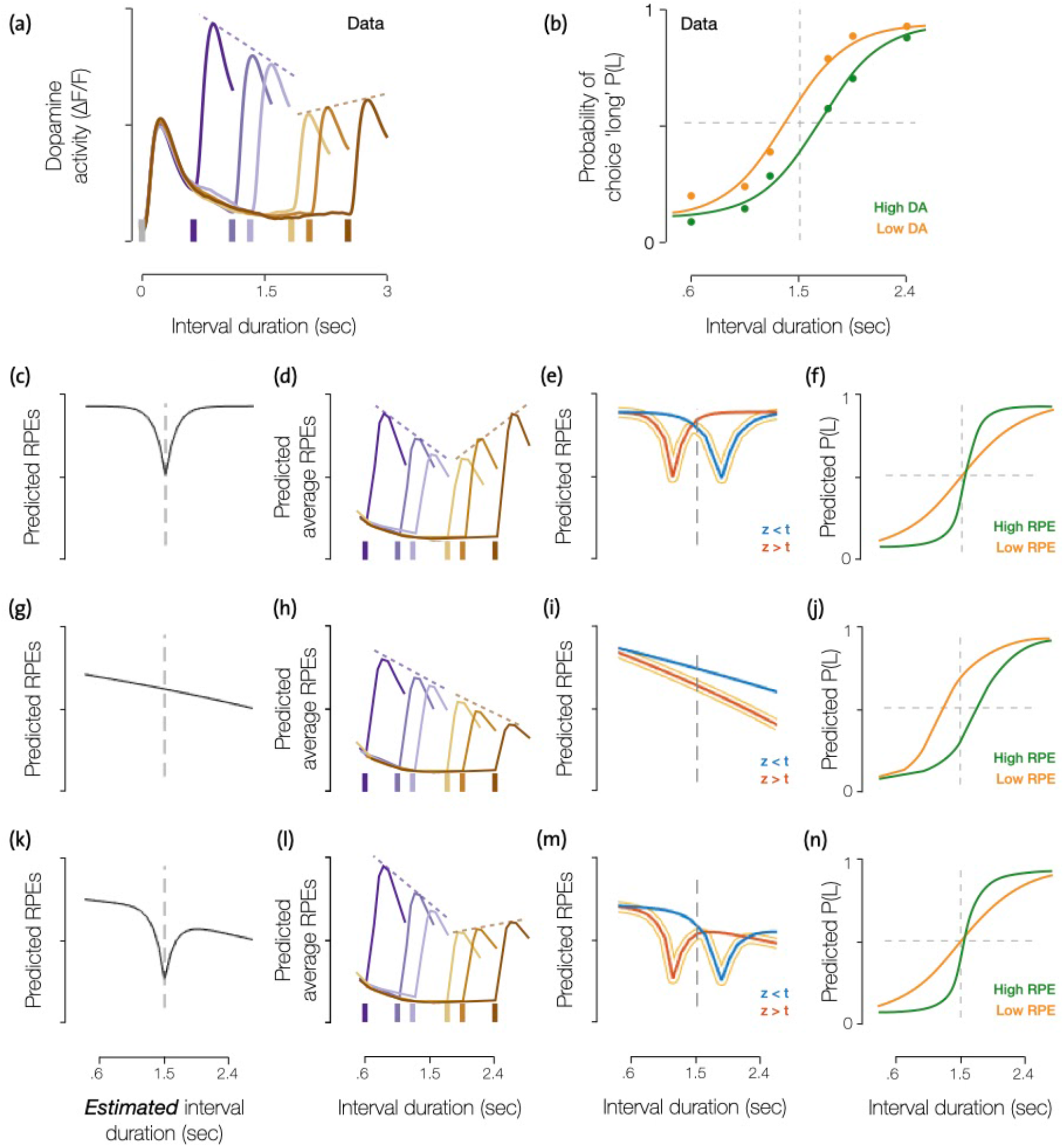
Reward prediction errors at interval offset can be modulated by choice accuracy and hazard rate of interval offset. **(a)** Average dopamine responses for each of the intervals presented during the task. The initial peak occurs at interval onset, and the following six peaks occur at each of the presented interval offsets. The dashed lines highlight the overall profile of the magnitude of responses at interval offset. **(b)** Psychometric functions for all trials in which high (green) or low (orange) DA responses were measured at interval offset. A clear difference in bias emerges from these two groups of trials. (Adapted from Soares et al 2016.) **(c)** If we assume that animals do not maintain any prediction of the arrival times of interval offsets, but do encode estimates of choice accuracy for different estimates of elapsed time, the RPEs we would predict would vary as a function of their internal estimates of elapsed time as shown here (For details see Supp. Fig. 1a-d). **(d)** Hypothetical modulation of average RPEs at the six interval offsets presented in the task if these were based purely on estimates of an agent’s choice accuracy. The averages in this case would be over the trial to trial variability in animals’ estimates of elapsed time. The dashed lines highlight that, unlike in the data, the overall profile of magnitude of RPE at interval offset would be symmetric around the decision boundary in this case. **(e)** Since animals’ reward expectations evolve as a function of their internal estimates of elapsed time, shown here is how their estimates would evolve as a function of real time on two example trial types where the animal may overestimate (red) or underestimate (blue) elapsed time. For any interval, whether interval duration is over or underestimated will systematically influence the magnitude of RPE at interval offset. Hence, for every time step, the curve that corresponds to low RPE is highlighted in yellow. **(f)** For every interval offset presented, if trials are split based on the magnitude of RPE, we would find that high RPE would go along with estimates of interval duration that are further than the boundary than those trials on which RPE is lower. Hence, when trials are split into low and high magnitude RPE trials, the slope of the psychometric curves of the two groups would differ. **(g-j)** Similar to (c-f), but for RPEs that are generated entirely due to temporal predictability of interval offsets. **(k-n)** Similar to (c-f) and (g-j), but for RPEs that would be generated if the agent took into account both choice accuracy and temporal predictability of interval offsets. For more details regarding the shape of the predicted RPEs in (c,g,k), see Sup. Fig. 1 and for more details regarding the predicted relationship between single trial magnitude of RPE and choice shown in (f,j,n) see Supp. Fig. 2.

After interval onset, the task required animals to maintain an ongoing estimate of elapsed time to guide their decisions. This internal estimate may be used not only to guide decisions, but also to encode time-varying expectations of future rewards. During the interval, expectations of future rewards should reflect average rewards expected from all intervals that are still possible. At interval offset, however, animals’ reward expectations should change and reflect an estimate of average rewards only from the estimated interval duration. Hence, when interval offset is presented, the change in reward expectations should cause RPEs. Since reward expectations, both before and after interval offset, depend on interval duration we expected DA responses at interval offset to also vary as a function of the duration of the interval. Indeed, we found that the magnitude of average dopamine response at interval offset was modulated by interval duration (Figure 2a). More specifically, two trends stood out. First, DA responses are larger for intervals further away from the decision boundary than for those close to the boundary. Second, the overall magnitude of responses is lower for ‘long’ intervals compared to ‘short’ intervals. Moreover, trial-to-trial variability in magnitude of DA responses also has a systematic relationship with animals’ reported judgments. For each interval duration presented, if trials are split based on the magnitude of DA response at interval offset into low and high magnitude trials, the psychometric function of trials that correspond to higher response magnitude is shifted right relative to that corresponding to trials with lower response magnitude at interval offset (Figure 2b). In other words, trial-to-trial variability in magnitude of DA response for each stimulus is predictive of the ‘bias’ in duration judgments. We will now investigate these trends in more detail.

At interval offset, since animals have not yet received any reward feedback, differences in RPEs and DA responses must be based only on internal variables. Previous work has shown that, when the timing of a sensory cue in the a task is variable, animals do estimate the hazard rate of the cue, i.e., the probability of the cue occurring at a given time given that it has not occurred yet (Janssen & Shadlen, 2005). Animals have been shown to encode their own choice accuracy as a function of stimuli presented, i.e., animals’ estimate their own ability to correctly classify different stimuli (Kepecs et al., 2008; Kiani & Shadlen, 2009). Separate studies have also shown that DA activity reflects changes in reward expectations based on the hazard rate of cues and rewards (Fiorillo et al., 2008; Pasquereau & Turner, 2014; Starkweather et al., 2017), as well as by animals’ choice accuracy (Lak et al., 2017). Hence, we expect animals’ reward expectations in the interval discrimination task to be influenced by hazard rate of interval offset as well as choice accuracy associated with each interval. To understand their joint influence, we first consider separately how choice accuracy and the hazard rate of interval offset might influence reward expectations and the consequent RPEs in the interval discrimination task.

### Choice accuracy may modulate RPE

Due to trial-to-trial variability in estimates of elapsed time, animals correctly categorize intervals close to the decision boundary less often than intervals far away from the boundary (Figure 1b). If animals keep track of the resulting choice accuracy as a function of interval duration, then they will expect less reward close to the boundary and more reward far away from it. If animals do not have any time-varying reward expectations during the interval, RPEs at interval offset will be modulated only by differences in reward expectation due to differences in choice accuracy as shown in Figure 2c (for more details see Supp. Fig. 1a-d). In other words, RPEs should be lower for estimates of interval duration close to the boundary than those further away. If we assume that animals’ reward expectations during the task are driven only by estimates of choice accuracy, then we should expect average DA activity to be modulated as shown in Figure 2d.

Moreover, animals’ errors in estimating elapsed time should also influence RPEs on a trial by trial basis. When animals over- or underestimate elapsed time, their internal estimates of reward expectations likewise evolve faster or slower in time (as shown in Fig. 2e). As a consequence, their internal estimate of choice accuracy and probability with which they will choose long’ or short’ will shrink and stretch with elapsed time in the interval. For any given interval duration, we should also find that individual trials with high RPE correspond to trials on which animals’ estimates of interval duration are further away from the boundary and hence are associated with high choice accuracy and low choice variability. In turn, trials with low RPE correspond to trials with low choice accuracy and hence high choice variability. Consequently, if we group trials based on the magnitude of RPEs at interval offset and plot the psychometric functions for the two groups of trials, the trials with higher RPEs will have a psychometric function with a steeper slope than those for low RPE trials (Figure 2f, for more details see Supp. Fig. 2a-d).

### Predictability of interval offset may modulate RPE

Similarly, we can consider the influence of the hazard rate of interval offset. On each trial, the interval is randomly drawn from one of six intervals, with equal probability. The hazard rate is thus a monotonically increasing function. In other words, the probability of encountering an interval offset at any time during the interval increases with time in the interval. If reward expectations are based only on hazard rate of interval offset, they should also increase monotonically with time in interval. When an interval offset is presented, the change in reward expectations will only reflect the fact that the presentation of the interval offset removes the uncertainty in the timing of the reward. As a consequence, the resulting RPEs at interval offsets later in the interval should be lower than those earlier in the interval (as shown in Fig 2g, for more details see Supp Fig 1e-h). If animals only encode the distribution of interval durations but not the outcome of their own choices, average RPEs elicited at interval offset will be on average lower for longer intervals than shorter ones, and we should expect the modulation of average DA activity shown in Figure 2h.

Due to trial to trial variability in estimating elapsed time, animals’ estimates of the hazard rate of interval offset and hence reward expectations will also vary from trial to trial (as shown in Fig. 2i). However, in this case, we will find that for any given interval offset, trials with high RPEs correspond to trials on which animals underestimate elapsed time, irrespective of the time at which interval offsets are presented. Consequently, if we group trials based on the magnitude of RPEs for each interval presented, the psychometric functions of the high and low RPE trials will show a horizontal shift, or, in other words, a change in bias (Figure 2j, for more details see Supp. Fig. 2e-h).

### Dopamine activity and its relation to choices can not be predicted by directly combining choice accuracy and predictability of interval offset in time

Finally, we consider the case when reward expectations are modulated by both choice accuracy as well as the hazard rate of interval offset. Since we assumed that choices are reported as soon as interval offset is presented, reward expectations at interval offset are the same as in the case when reward expectations are modulated only by choice accuracy. However, during the interval, reward expectations should reflect the fact that, both, the timing of interval offset is unpredictable but also that different interval offsets predict reward with different probabilities based on animals’ choice accuracy. Hence RPEs in this case will reflect both choice accuracy as well as the hazard rate of interval offset as shown in Fig. 2k (for more details see Supp. Fig. 1i-l). In this case, we expect that average RPEs at each interval offset may reflect both choice accuracy as well as the hazard rate in a manner that may be similar to that in the data (as shown in Fig. 2l).

Again, trial to trial variability in estimates of elapsed time will affect the time evolution of reward expectations (as shown in Fig. 2m) and choice probability. In this case, for any given interval offset, we find that high RPE trials will correspond to trials on which the agent’s estimate of the duration is further from the boundary and hence will be associated with higher choice accuracy and low choice variability. Consequently if group trials, for each interval presented, based on the magnitude of RPE on that trial, we see a difference in the slope of the psychometric curve of the two groups of trials (as shown in Fig 2n, for more details see Supp. Fig. 2i-j).

In other words, animals’ choice accuracy, the hazard rate and the combination of the two predict distinct patterns of RPEs at interval offset. The two key experimentally observed trends--the profile of average DA responses at interval offset and the trial to trial relationship between magnitude of DA for any single interval offset and animals’ choices--can not be simultaneously explained by either of these three strategies. Average DA responses are best captured by computing reward expectations that take choice accuracy as well as hazard rate of interval offset into account (Figure 2a is consistent with 2l). However, the differences in the psychometric functions for high and low DA is captured by computing reward expectations that only take into account the hazard rate of interval offset (Figure 2b is consistent with 2j).

Since our simplified considerations of how RPEs (and thereby DA responses) should be affected by temporal predictability and choice accuracy do not match the data fully, we hypothesised that our assumptions regarding how task variables may be encoded by animals, based on which we computed reward expectations, may not be accurate. To gain a more detailed understanding of how differences in the encoding of task variables as well as animals’ own behaviour may influence RPEs, we simulated a reinforcement learning agent that could make choices at any time during the trial and was required to learn from trial and error, just like animals, to take the right action at every timestep to obtain rewards.

### RL agents were modeled to keep track of time since task events

Since animals’ choices are based on variable internal internal estimates of elapsed time, we modelled the reinforcement learning agent using a partially observable Markov decision process (for details see Methods section 1). We assumed that the agent keeps track of the timing of events in the task within each trial and we allowed the agent to make choices at any time during the trial. More specifically, we assumed that the agent can maintain noisy estimates of elapsed time since interval onset and elapsed time since interval offset. With these two estimates, the agent has all the necessary information to estimate the length of the interval presented as well as the task epoch, which in turn should inform the agent’s actions at all other time points in the task. The agent needs to learn from experience to withhold choices during the interval and to report choices based on its estimate of interval duration after interval offset. Figure 3 shows the state space encoded by the agent and how it traverses through the state space on two example trials types. In both examples, the agent encodes elapsed time since interval onset by advancing horizontally in the depicted state space. After interval offset, the agent additionally encodes elapsed time since interval offset and therefore traverses the state space along diagonals. Since time since interval onset will always be larger than time since interval offset, the agent will only visit states that are below the unity line in this state space. For all states encountered, the agent needs to learn the optimal action to take. During the interval it needs to learn to withhold choice and after interval offset the optimal action depends on the x-intercept of these diagonals, i.e., on the difference between its estimates of elapsed time since interval onset and interval offset.

**Figure 3:**
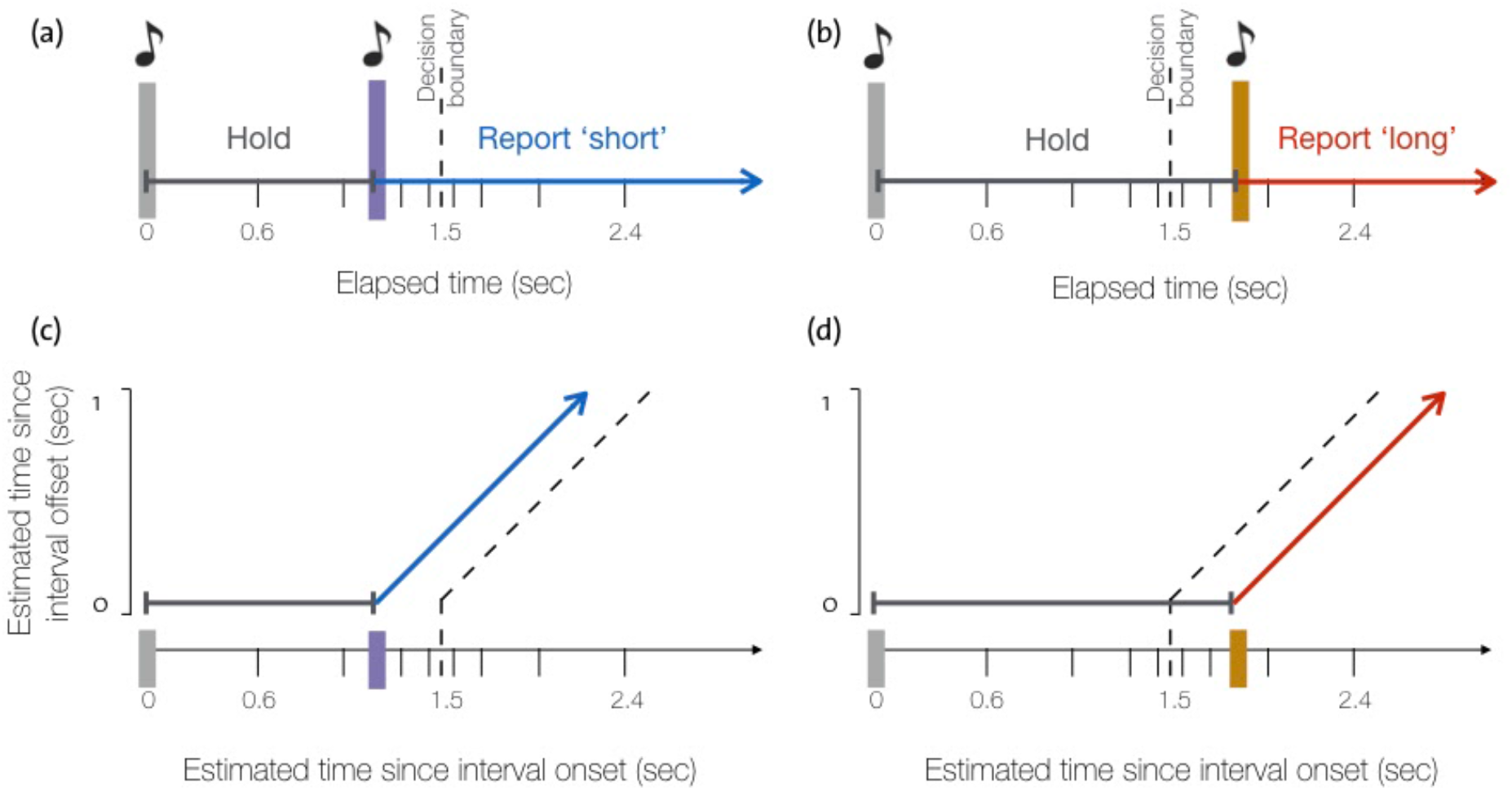
Task variables can be represented unambiguously using a two-dimensional state space. **(a)** Timeline of an example trial in which an interval shorter than the decision boundary is presented. The color of the axis indicates the optimal action in that period of the trial (grey = wait, blue = short, red = long). **(b)** Timeline of an example trial in which an interval longer than the decision boundary is presented **(c)** Illustration showing how the example trial in (a) is represented in the agent’s state space. The agent’s state is given by its internal estimates of time since interval onset (x-axis) and time since interval offset (y-axis). Because of variability in time estimation, these internal estimates are distinct from real time. The trajectory shows how the two-dimensional state variable changes over time and in response to interval onset and offset. The color of the arrows shows the correspondence between the segments of the trajectory in state space and the associated segments on the timeline of the example trial in (a). Each location in the two-dimensional state space provides an unambiguous representation so that the agent can determine the optimal action. The internal decision boundary that would allow the agent to make optimal choices using this state representation is shown by the dashed line. **(d)** Illustration showing how the example trial in (b) would be represented in the agent’s state space.

The agent learns using TD-learning within an actor-critic architecture (for more details, see Methods). Actor-critic architectures have been commonly used to model dopamine activity as RPEs in tasks where outcomes depend on actions taken by animals (Joel et al., 2002; Khamassi et al., 2005). In this framework, the agent estimates reward expectations from each state (i.e. state-value function) and a state-action mapping (i.e. policy), that instructs the agent which action to take, for all states it encounters. The agent learns to select actions that result in transitions to higher-value states and hence those transitions become more probable than those that lead to states with low value. Importantly, the state value function and policy must both be learned simultaneously and are both defined as functions of the location in state space. Since the agent’s internal estimates of time are modeled as continuous variables, there are infinitely many locations in state space the agent could be in. This makes the task of learning value functions for each state directly highly impractical. Hence, we use a function approximation scheme to estimate the value function and policy. Both these functions are approximated using a set of basis functions or feature vectors (Figure 4a shows a schematic of the function approximation scheme used). To keep this approximation as simple as possible, we used non-overlapping tile bases. This is equivalent to discretizing the continuous state space for value approximation.

**Figure 4:**
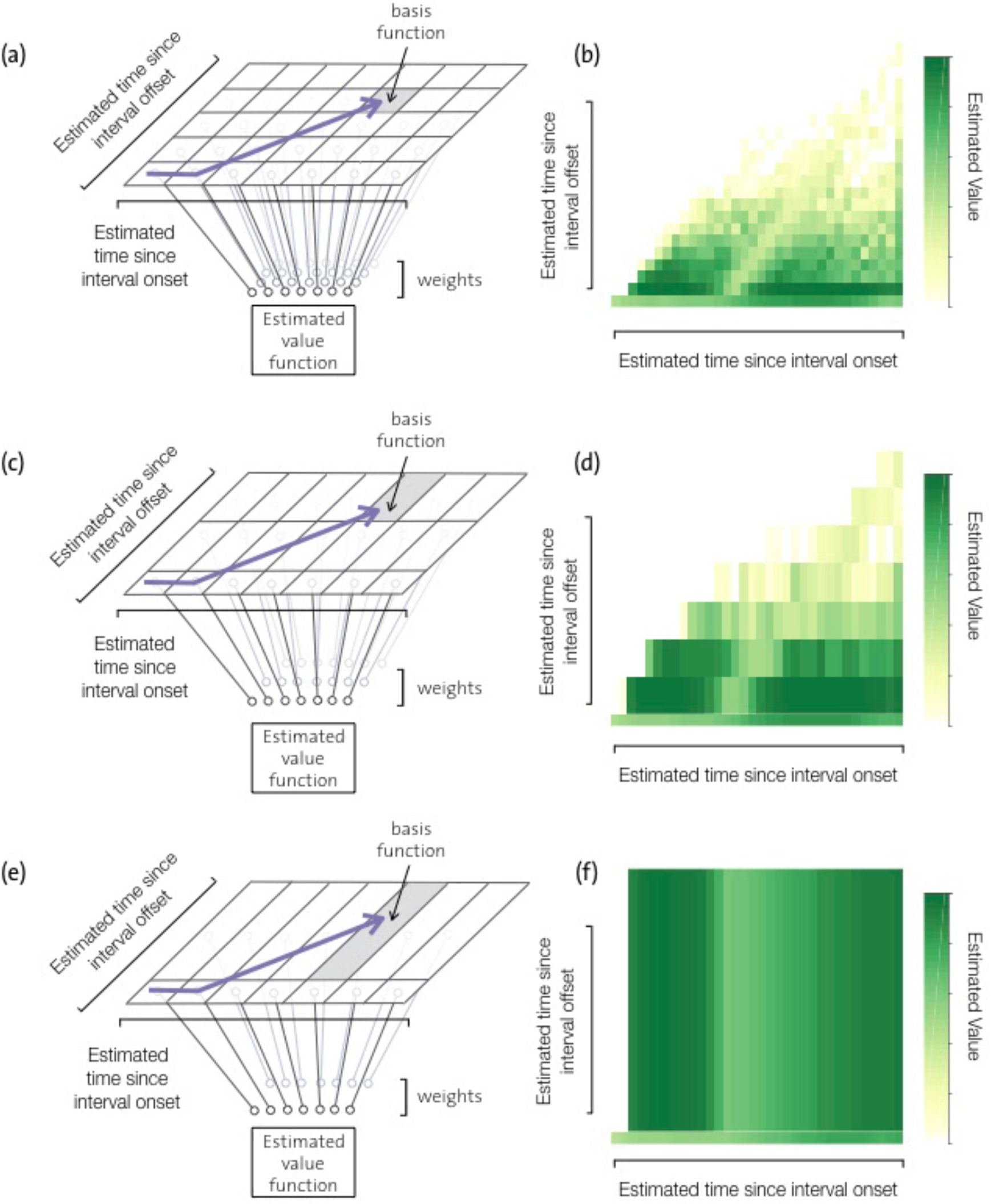
Resolution of the basis functions used to approximate value function and policy can be different along different dimensions of state space. **(a)** Since the state space is continuous, we approximate the value function by a linear combination of basis functions that are non-overlapping tile bases. The agent needs to estimate as many parameters or weights as there are basis functions. Here the tiling of the basis functions is equally dense along the two axes shown and hence the number of parameters that need to be estimated are ~N^2^, where N is the number of basis functions tiling each dimension. This arrangement corresponds to the full mapping used by the model. Shown in purple is an example trajectory through the state space. The basis function that would be active at the last time step of this trajectory is highlighted. **(b)** Estimated values learnt over the state space using the full mapping. **(c)** Basis functions used by the agent can tile the second axis with lower resolution than the first, thereby incurring a lower representational cost than the full mapping. This example corresponds to an intermediate level of compression. **(d)** Estimated value function using the intermediate compressed mapping. **(e)** The basis functions used in the most compressed or efficient mapping tile the second axis with the lowest possible resolution by encoding only whether interval offset has occurred or not. The number of parameters that need to be estimated using this mapping are ~2N (as opposed to ~N^2^ parameters that need to be estimated for the full model). **(f)** Estimated value function learnt using the most compressed, i.e. efficient representation.

Previous experimental findings have shown that animals exhibit trial-to-trial variability in estimating elapsed time and that the standard deviation of variability in timing estimates increases linearly with time, which is known as the scalar property in timing (Gibbon & Church, 1990). Hence, we constructed the agent’s internal representation to also have trial-to-trial variability in estimates of elapsed time that obeys this scalar property (for more details see Methods). The amplitude of the noise was adjusted so as to qualitatively match animals’ overall task performance (determined using the psychometric function).

### RL agent constructed based only on task requirements cannot reproduce relationship between DA response and choices

We first modeled an RL agent that estimates the value function and policy by uniformly tiling the state space along both dimensions of the state space (Figure 4a, 5a). After the agent learns the task, we find that the profile of average RPEs at each interval offset qualitatively captures that of average DA activity (compare Figure 5b with Figure 2a). However, the trial to trial relationship between the magnitude of RPEs and the agent’s choices is inconsistent with what we see in the data (compare Figure 5c with Figure 2b). Rather, the psychometric functions of the high RPE and low RPE groups of trials show a change in bias as well as slope, as seen when we directly computed reward expectations based on choice accuracy and predictability of interval offset in time (see Fig 2n and Supp. Fig. 2i j). n other words, the trial to trial relationship of the agent’s RPEs with its temporal judgements is influenced by discrimination accuracy as well as hazard rate of interval offset. At first sight, these results suggest that there might be some aspect of dopamine activity that cannot be captured entirely by RPEs during this task. However, since RPEs are calculated from the agent’s expectations of future rewards, given by the state value function (Figure 4a), the results could also suggest that animals are calculating expectations of future rewards in a way that does not match the true underlying structure of the task. Such a mismatch could come about if the representation that the animals are operating on are misrepresenting the statistical structure of the task.

**Figure 5:**
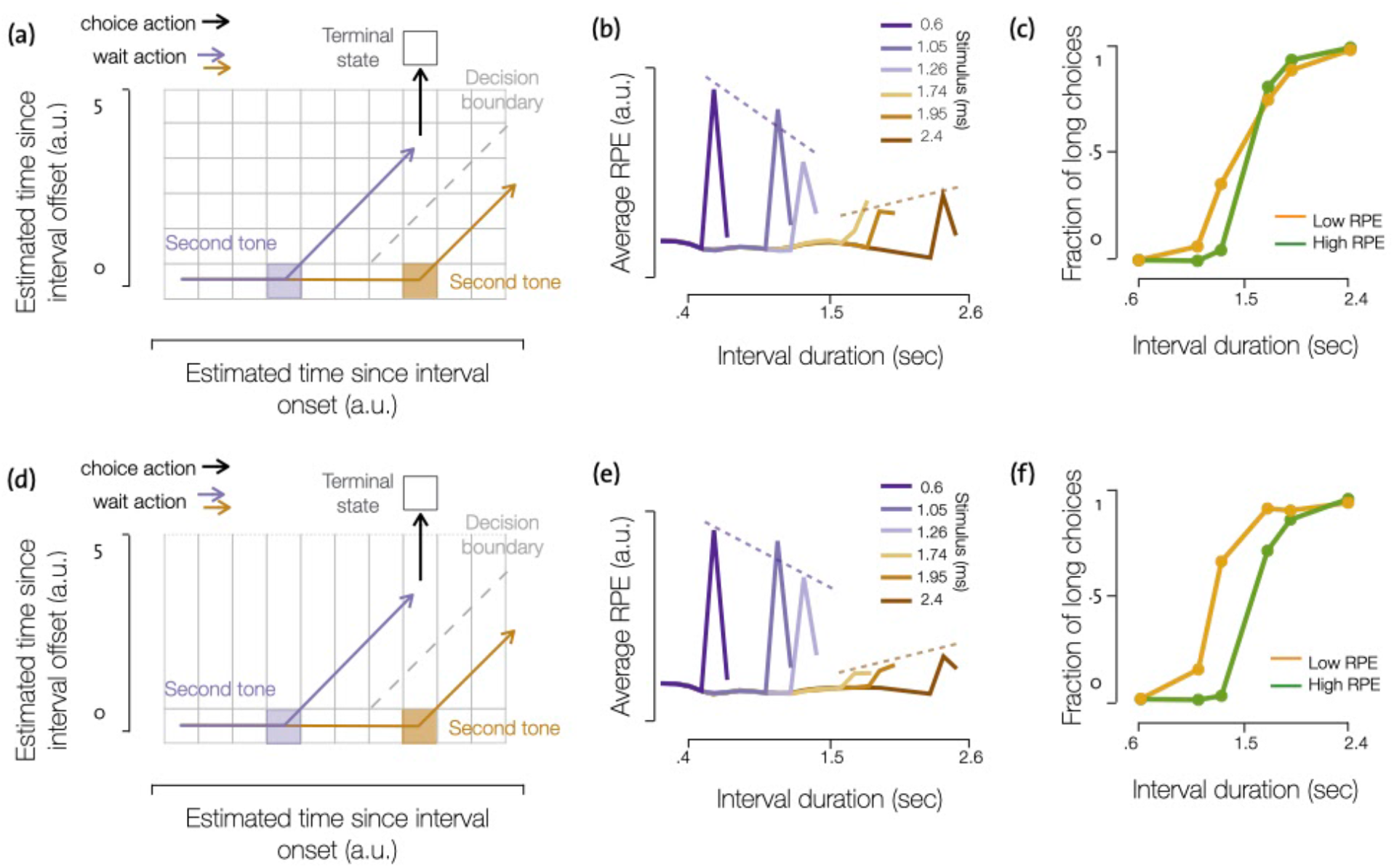
Efficient value function approximation can simultaneously reproduce average DA at interval offset and trial to trial relationship between DA magnitude and choice. **(a)** Value funct on approx mat on used n the full mapping. Each basis uniquely determines est mated elapsed time since interval onset and interval offset. **(b)** Average reward prediction errors (RPE) elicited at interval offset n an agent that uses the full mapping for interval discrimination. Compare with Figure 2a. **(c)** Psychometric curves of trials grouped based on the magnitude of RPE at each interval offset for an agent us ng the full mapping. Compare with Figure 2b. **(d)** Value function approximation used in the efficient mapping. **(e)** Average reward prediction errors (RPE) elicited at interval offset in an agent that uses the efficient mapping for interval discrimination. Compare with Figure 2a. **(f)** Psychometric curves of trials grouped based on the magnitude of RPE at each interval offset. Compare with Figure 2b.

To ask how the underlying state representation could be different, we note that the dynamics of the latent variable in the task is not made available to animals and that they need to infer how the task should be represented using only the sparse observations they recieve. Previous work has shown that population dynamics in the striatum during the interval encode elapsed time with high fidelity and that trial to trial variability in how activity evolves is predictive of animals’ temporal judgements (Gouvêa et al., 2015). Hence, we maintained high resolution with which time since interval onset was encoded in the model. Other work, in the context of an interval reproduction task, has shown that cortical dynamics encode elapsed time since interval onset and offset in a similar manner (Remington et al., 2018). Based on these findings, we assumed in the model implementation above, that animals would represent elapsed time since interval onset and offset in a similar manner, with high fidelity, during an interval discrimination task as well. However, this assumption may not be well aligned with how animals may be representing the interval discrimination task. Drawing on the principles of efficient coding, we reasoned that in addition to maximising the overall number of rewards obtained during the task, animals may want to minimise the computational resources required to estimate the value function and policy over all possible states. n particular, we hypothesised that animals’ may encode time varying value functions with higher resolution during the interval and but not after the interval. Hence, we asked whether and how encoding elapsed time since interval onset and offset with different resolutions would affect task performance as well as reward expectations learnt during the task.

### RL agent with efficient representation can reproduce trial-to-trial relationship between DA response and choices

Let us consider the function approximation used to estimate the value function described in Figure 4a. The size of each basis function (or tile) will determine how accurately changes in value from one location to another can be estimated. In turn, the number of basis functions (or tiles) will determine the computational cost of estimating the entire value function. Accordingly, more accuracy incurs more costs. An efficient coding scheme might stipulate that the resolution with which an optimal set of basis functions tile the two dimensions of the state space should depend on the degree to which the readout varies along each of these dimensions (Salinas, 2006). More specifically, value functions along the axis that represents estimates of elapsed time since interval onset may be encoded with high resolution, since doing so is necessary for the agent to accurately report choices. Value functions along the axis that represents estimates of elapsed time since interval offset, on the other hand, may be encoded with a lower resolution, since lack of accuracy here may not adversely affect the animal’s ability to make the correct choice (see Figure 4c,e). In the extreme case, when the axis representing time since interval offset is encoded with the lowest possible resolution (while still encoding interval offset), the basis functions effectively encode only time since interval onset and whether or not interval offset has occurred (Figure 4e). We will refer to the basis functions as described in Figure 4a as the high resolution or full mapping and the other extreme shown in Figure 4e as the low resolution or representationally efficient mapping.

When we train the RL agent using the efficient mapping, we find that it is able to obtain similar numbers of rewards as an agent trained using the full resolution mapping. The efficient model also reproduces the profile of average DA responses at interval offset (compare Figure 5e and Figure 2a). In other words, the compression of the mapping along the second axis did not adversely affect the agent’s choice behavior, nor did it change the predictions for average RPEs at interval offset. Surprisingly, however, the efficient representation is also able to reproduce the trial to trial relationship between magnitude of DA and temporal judgements (compare Figure 5f and Figure 2b). This somewhat puzzling result was only obtained for very strong compressions of mappings along the second axis and not for intermediate levels of compression. When we simulated the agent using several intermediate levels of compression in representing elapsed time since interval offset, we obtained results more similar to the unconstrained agent (see Supplementary Figures 7 and 2). Accordingly, only a large difference in the resolution with which elapsed time since interval onset and offset are encoded can explain the observed DA responses and their relation to behavior.

We would like to note that we have focused primarily on describing how changing the basis functions, without changing the dynamics of the latent variable, can achieve a more coarse approximation of the value function along the second dimension of the latent variable. However, we could have achieved the same result by keeping the basis functions the same and changing the dynamics of the latent variable. Since the nature of value approximation only depends only on the relationship between the latent variable and the basis functions, for simplicity we only explain in detail our implementation that changes the basis functions and not dynamics of the latent variable (for more details see Methods section 4). We would also like to note that, although the efficient mapping that we consider in detail is as shown in Figure 4e, there are indeed multiple ways to specify the dynamics of the latent variable or the mapping to approximate value functions in our 2D state space that would be equivalent to the efficient representation we discuss in terms of number of parameters to be estimated for value approximation. However, none of these alternatives allow us to reproduce the data shown in Figures 2a and 2b simultaneously. Only when the relationship between the latent variable and the mapping is kept the same as in the efficient representation we discussed, can the RL model reproduce the observed data. Some of these alternatives are discussed in Methods section 5.

### Reward expectations at interval offset are markedly different under the different mappings

To understand why our proposed efficient mapping results in very different RPEs, we look at the value function learned by the agents using the full vs efficient mappings (as shown in the schematics in Figure 4a and 4e respectively). We note that RPEs at interval offset reflect the difference between the agent’s reward expectation (given by the corresponding value function) during the interval and its reward expectation at interval offset, at which point the agent does have an estimate of the interval duration to be classified on that trial. For agents using the full mapping, reward expectations before interval offset reflect the hazard rate of interval offset and choice accuracy, which increase as a function of elapsed time (Figure 6a). After interval offset, the agent has acquired an estimate of the presented interval, and, therefore, its reward expectations only reflect choice accuracy (Figure 6b). At any time, the difference between these two value functions determines the RPE if interval offset was presented at that time (Figure 6c). Consequently, RPEs at interval offset reflect both the hazard rate of interval offset and the agent’s choice accuracy.

**Figure 6:**
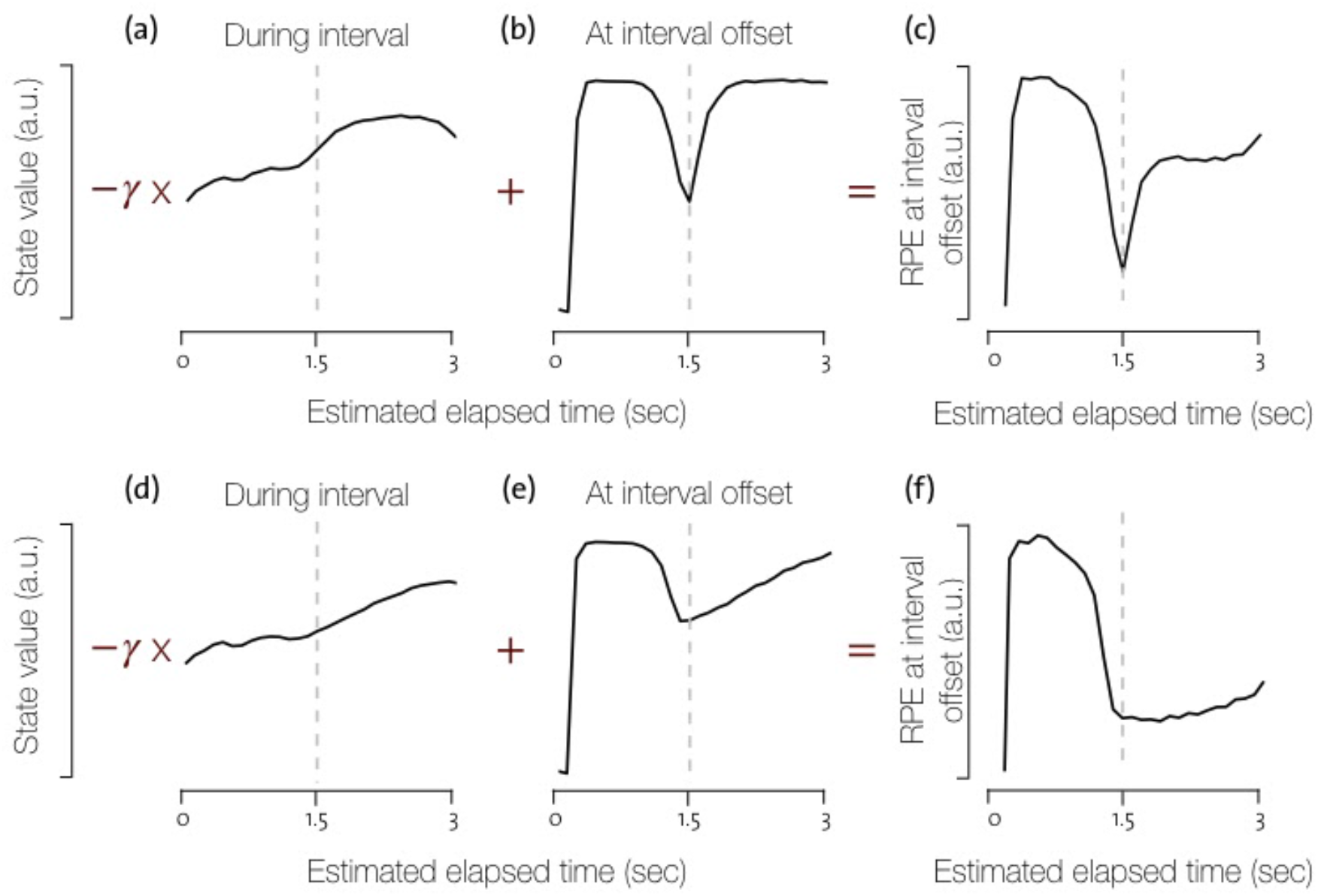
Reward expectations learnt by the RL agent using the full and efficient mappings differ most right after the decision boundary. **(a-b)** Reward expectation during the interval (a) and after interval offset (b) as a function of internal estimates of elapsed time since interval onset using the full mapping for value estimation. **(c)** Reward prediction error at interval offset as a function of all estimates of elapsed time since the interval onset using the full mapping. **(d-e)** Reward expectation during the interval (d) and after interval offset (e) as a function of internal estimates of elapsed time since interval onset using the compressed mapping for value estimation. **(f)** Reward prediction error at interval offset as a function of all estimates of elapsed time since the interval onset using the compressed mapping.

Similarly, we can look at the value function learned using the efficient mapping. As before, we see that reward expectations during the interval increase with the length of the interval (Figure 6d). However, reward expectations after interval offset do not simply reflect the agent’s choice accuracy. Instead, they exhibit a strong asymmetry around the decision boundary (Figure 6e). We see that on the long side of the boundary, reward expectations increase much more slowly as a function of distance from the boundary than on the short side of the boundary. Consequently, the resulting RPEs reflect this asymmetry and show a slower rise on the long side of the decision boundary compared to the short side (Figure 6f). Before exploring the origin of this asymmetry (discussed in the following section), we will show that it substantially changes the trial to trial relationship between the animals’ behaviour and dopamine activity.

Let us first consider two near-boundary intervals. For each of these intervals, the agent’s estimate of elapsed time will vary from trial to trial as described by the two distributions shown in Figure 7a and Figure 7d. We recall that the agent’s decisions are based entirely on its internal estimates of elapsed time: when the estimate is shorter (or longer) than the boundary, the agent will report choice ‘short’ (or ‘long’) with higher probability. We can therefore split each of the distributions based on the choice of the agent (Figure 7a,d, red and blue areas). This procedure creates four groups of trials, given by the two near-boundary intervals and the two choices of the agent. Within each group, the variability in time estimates gives rise to associated variability in RPEs. The four resulting distributions of RPEs are shown in Figure 7b,e. Here, the two panels group trials according to the presented interval, and the red and blue RPE distributions in each panel correspond to trials grouped according to the animal’s choice. Using these groupings, we can now study how high or low RPE trials relate to behavior. To do so, we first define high (or low) RPE trials for a given interval as all trials with RPEs greater (or smaller) than the median RPE for that interval (indicated by the dashed line). The fraction of long choices falling into the high (or low) RPE trials shown in Figure 7c,f correspond to points at the near boundary intervals in the psychometric functions for shown in Figure 5c (which are for all interval durations).

**Figure 7:**
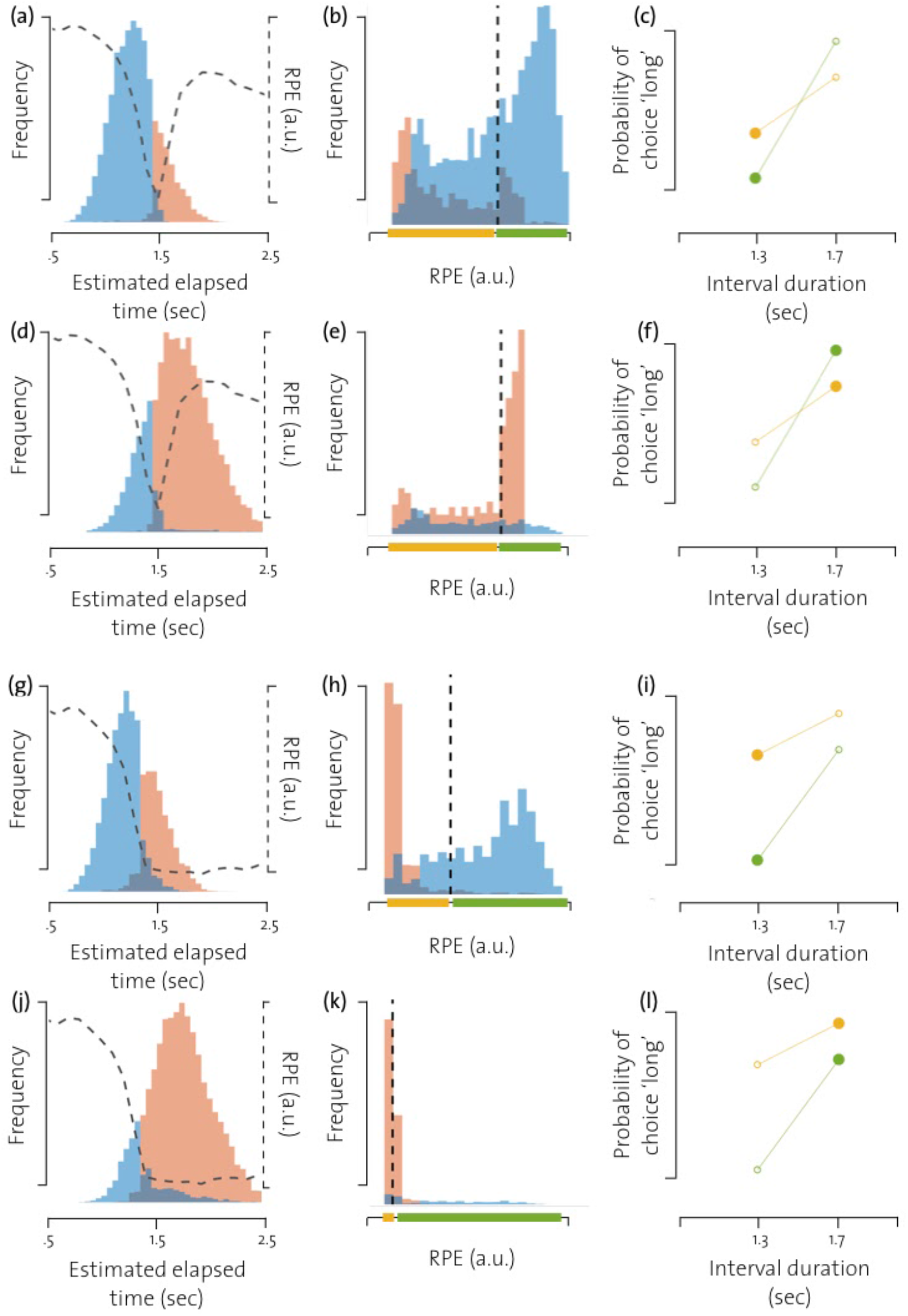
Distributions of RPEs at interval offset show why compression in the mapping causes psychometric curves split by RPE magnitude to be different. **(a)** Distributions of the agent’s internal estimates of elapsed time at interval offset for the ‘short’ near-boundary interval. The distribution is color-coded according to the agent’s choice (blue=short red=long). The dotted line shows RPEs at interval offsets for the corresponding estimates of elapsed time. (**b)** Distributions of RPEs at interval offset for the ‘short’ near-boundary interva. The distributions are colored according to the agent’s choice on the corresponding trials. The dotted line indicates median-RPE for that interval presentation and divides RPEs into a ow-RPE group (yellow bar) and a high-RPE group (green bar). **(c)** Fraction of trials on which the agent chose ong’ on trials with RPEs ower (yellow) or higher (green) than the median RPE for the two near-boundary intervals. Larger dots indicate the fraction of ong’ choices for the short’ interva. **(d-f)** As in (a-c), but for a ong near-boundary interva. **(g-l)** The same as in (a-f), using the efficient mapping.

For the RL agent using the full mapping, RPEs on incorrect trials are on average lower than RPEs on correct trials (Figure 7b,e). n turn, if we split all trials based on the magnitude of RPEs, irrespective of the interval, we find that the agent made more mistakes on low RPE trials compared to high RPE trials. Thus, the psychometric curves for these two groups of trials show a larger difference in slope and cross each other around the decision boundary (Figure 7c,f). For the RL agent using the elft cient mapping, the picture is very different (Figure 7g-l). Here we find that RPEs are on average lower when the agent reports choices as long, irrespective of the interval (Figure 7h,k). Consequently, the psychometric functions for high and low RPEs show a larger change in bias and that the psychometric curves for these groups of trials do not cross each other near the boundary (Figure 7j,l).

### Efficient mapping predicts procrastination of choices for interval durations estimated as long and close to the decision boundary

Finally, we studied the origin of the asymmetry in the value function for the agent using the elft cient mapping. We find that the low-resolution along the second axis leads to a systematic ambiguity in some parts of the state space in estimating value and the optimal action for those locations. For example, let us consider a basis function that spans a region in state space that the agent would visit right after the end of a long interval (marked with the gray rectangle in Figure 8a). While the agent can encounter this region directly after the end of a long interval, it could also be encountered if the agent was presented with a short interval, but withheld choice for several time steps (purple trajectory in Figure 8a). Hence, the reward expectation associated with this basis will be estimated by averaging over the trials from both categories of interval durations i.e. when the agent’s estimate of the interval presented is longer or shorter than the learnt boundary. As a consequence, the agent would be impeded in learning the correct value of these states as well as the optimal action at these locations. Indeed, this is true for all the post-interval basis functions in the efficient mapping. They encode elapsed time since interval onset and whether or not interval offset had occurred. Accordingly, the basis functions do not allow the agent to disambiguate between trials on which different interval durations were presented if it withholds choice for several time steps.

**Figure 8:**
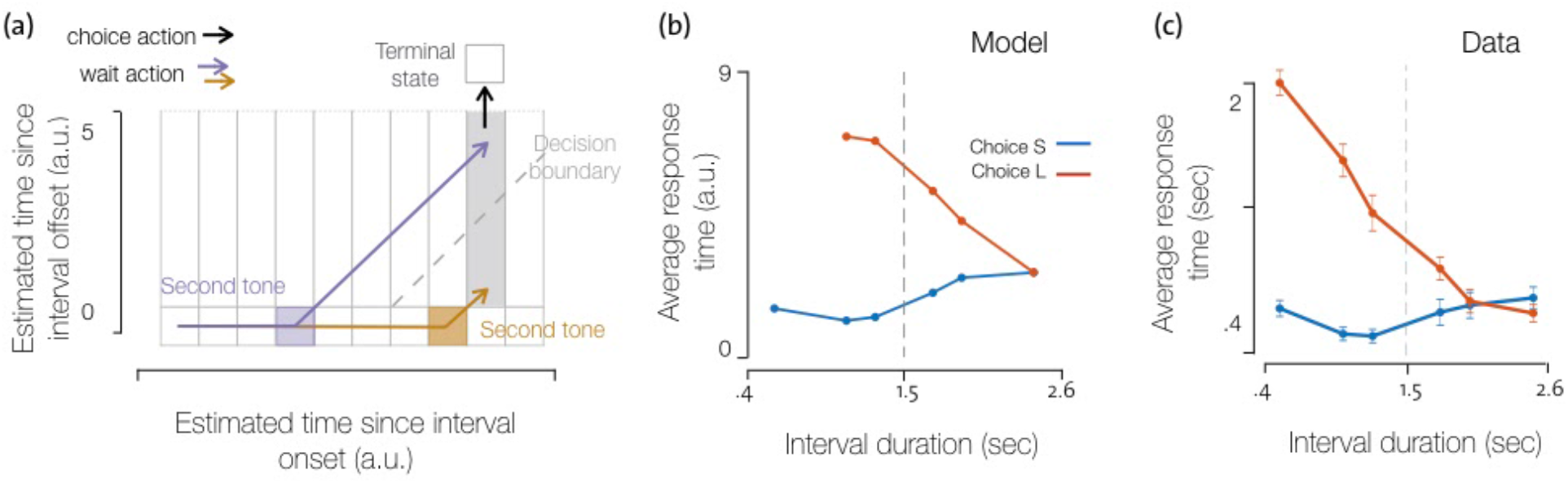
The efficient mapping predicts an unusual profile of response times that closely matches animals’ behaviour. **(a)** Efficient mapping and two example trajectories through the state space. The grey rectangle indicates a post-interval offset basis. Note that this region of the state space can be reached following long intervals (yellow) as well as short intervals (purple) if the agent waits and withholds choice. The basis function is therefore ambiguous in informing the agent which interval duration was presented when the corresponding states are encountered on any given trial. **(b)** Average response times of an RL agent that uses the efficient mapping, conditioned on whether they indicated the interval to be short or long with respect to the decision boundary. **(c)** Animals’ average response times after each interval offset split based on choice.

The ambiguity in this efficient mapping can be avoided if the agent reports choices immediately after interval offset, especially for intervals estimated to be shorter than the decision boundary. Once the choice is made, the agent transitions into the inter-trial interval state, and thereby avoids visiting other post-interval states. With this strategy, states that fall into the region indicated by the grey rectangle in Figure 8a are only encountered when the agent estimates the interval to be longer than the decision boundary. In turn, the agent can learn that the correct choice associated with those states is ‘long’. Indeed, we find that when agents use the efficient mapping and when they estimate the interval to be shorter than the decision boundary, they report choices with very short response times (Figure 8b). For intervals longer than the decision boundary, however, there is no urgency to respond after interval offset. In this case, if the agent waits after the interval offset, the correct action associated with the post-interval offset states will not change with the passage of time. Hence, delaying choice will not have any detrimental effect on choice behaviour. Curiously, however, the efficient mapping not only allows delaying near boundary long choices, but incentivises it.

To understand why, let’s consider the value function of an agent using the efficient mapping, but where the agent is forced to report a choice immediately after interval offset. In this case, the value function estimated for the post-interval offset states using the efficient model is the same as that of the states immediately after interval offset using the full-mapping. After interval offset, states further from the boundary have higher value than those close to the boundary. Now, let’s say, the agent is permitted to have variable response times. Let’s first consider the value of the sequence of states the agent would encounter if it received a ‘short’ interval and withheld choice after interval offset (shown by the purple trajectory in Supp. Fig. 5a and 5b). In this case, if the agent waits after interval offset, the value of the sequence of states encountered decreases with time (shown by the red arrow in Supp. Fig. 5b). Hence, if the agent withholds choices, it will encounter negative s as elapsed time advances i.e. it moves to states that have lower value than the preceding states. However, if it reports it’s choice immediately, it will, on average, get zero s since choices lead the agent to transition to the terminal state and obtain rewards which are on average equal to the value of the state just preceding the choice. Since, the policy of the RL agent is to choose actions that lead to states with the highest value, the optimal strategy to report choices immediately. On the other hand, if the agent receives a long’ interval (shown by the brown trajectory in Supp. Fig. 5a and 5c), especially one close to the decision boundary, we see that the value of the sequence of states after interval offset increases with time (shown by the red arrow in Supp. Fig. 5c). Hence, if the agent withholds choices, it will encounter positive RPEs as elapsed time advances i.e. it moves to states that have higher value than the preceding states. This is what drives procrastination of choices in the model when the presented interval is estimated to be long and close to the boundary. In turn, these transitions between states after interval offset flatten the value function of those states on the long side of the boundary and hence causes the asymmetry we see in the value function when using the efficient mapping. The interaction between the efficient mapping and the policy learned when using it, causes the RL agent to learn reward expectations that generate RPEs similar to the dopamine activity recorded in animals performing the task.

In several two-alternative decision making paradigms, animals generally take longer to respond when the decision variable is close to the decision boundary (Roitman & Shadlen, 2002). Several tasks in which response times are longer for harder stimuli are tasks which require integration of noisy evidence, where the noise is uncorrelated over time. In these contexts, longer response times allow animals to integrate over more samples of noisy evidence and have a better estimate of the stimulus by averaging out the noise. In other words, longer response times result from increased deliberation when the stimulus category is more ambiguous. However, the response times of the RL agent using the efficient mapping in our task are markedly different, as they predict long responses only for difficult long’ choices, but not for difficult short’ choices. In other words, the RL agent generates a highly non-trivial prediction that can be tested against data. The long RTs we see for intervals that are perceived by the agent to be near-boundary long’ appear to be a result of *procrastination* of difficult long’ choices. Surprisingly, we find that animals also procrastinated as predicted by the model. The predicted pattern of response times from the model closely resembles that of animals during the task (Figure 8c). We also find that this pattern of response times is not reproduced by the agent when using the full mapping. Moreover, the profile of response times is observed only for highly compressed mappings (Supplementary Figure 4) and not for intermediate levels of compression such as the one shown in Figure 4b

Furthermore, if we force the agent to not have variable response times, the profile of psychometric curves of trials grouped by magnitude of RPEs at interval offset in the agent matches those resulting from the full mapping and do not match those in the data. Thus, animals’ pattern of response times provides further evidence to suggest that animals may indeed be using a representation similar to that captured by the efficient mapping while solving this task.

## Discussion

Understanding the nature of representations used by animals for value-based decision making is crucial to further our understanding of how neural circuits subserve adaptive behavioral control. Characterising what reward expectations animals learn can inform us about how and what variables they are representing, which in turn can reveal the constraints and strategies with which animals might be inferring statistical regularities in their environments. Here we studied the computations underlying a rigorously controlled time dependent behavior in rodents. We focused on two aspects of experimental data collected during this behavior, the recorded activity of dopamine neurons and its trial to trial relation to animals’ choices. Using an RL framework, we investigated how varying internal representations, with which the agent was able to solve the task, varied RPEs encountered by the agent. By comparing such RPEs with recorded DA responses, we were able to infer the nature of internal representations animals might be using during this task.

In several sensory systems, the principle of representational efficiency has been used to characterise neural coding. In many of these systems, the nature of the variable that is to be represented, and hence subject to efficiency constraints, is usually well defined. However, in the case of RL, it is unclear which variables used for value based choices might be subject to constraints of representational efficiency in the brain (Botvinick, et. al., 2015). For example, we may want to enforce efficiency constraints in how the structure of the environment is represented (Botvinick, Niv, & Barto, 2009; Wimmer, Daw, & Shohamy, 2012). This approach may be considered to be the closest to that used for efficient coding in sensory systems. In this case, the problem can be stated as that of identifying statistical regularities or redundancies in the environment that can be leveraged to balance representational cost against the accuracy with which the statistics of the environment can be encoded. However, this is not the only approach that can be used for representational efficiency in RL. Representation constraints may be used to directly approximate optimal value functions (Foster & Dayan, 2002) or action spaces (Solway et al., 2014). In each of these cases, one has to define a space within which representations are subject to resource constraints as well as the quantity of interest that needs to be preserved, i.e. a loss function which needs to be optimised. A key challenge in understanding how principles of efficient coding are applicable to the reward system is to identify the space within which representational constraints may be expressed as well as the loss function that may be optimised.

The interval discrimination task requires animals to generate and operate upon structured, time-evolving internal estimates to guide decisions. By training a RL agent on this task to generate dynamically evolving patterned behaviour, just as is required of animals, we were able to test how differences in task representation affect behaviour, which in turn affect reward expectations during the task. We found that animals seem to encode the task using a representation that is compromised in accurately capturing the statistical regularities of the task environment. This suggests that although our results are consistent with representational constraints being expressed in the space of how environmental states may be represented, the quantity being preserved is not the accuracy with which the statistics of the environment can be best captured. Nor does this representation allow the agent to well approximate the optimal value function that would be found using an unambiguous representation of the environment. This suggests that neither is the optimal value function the quantity that is being preserved. However, the representation used allows the agent to learn actions that result in equivalent number overall rewards while being more compact than one that would allow an unambiguous representation of all states in the task. Importantly, we find that the behavioural strategy the RL agent uses while using the efficient representation is crucial to prevent ambiguities in the representation from affecting the accuracy of choices and hence the overall number of rewards that can be obtained. Furthermore, only when the behavioural strategy of the agent is allowed to be influenced entirely by maximizing RPEs using the efficient representation is the model able to capture the key features in the data. This suggests that the representations used by the agent may be found by maximising the same quantity used by the agent to select actions i.e. overall number of rewards (or by minising TD reward prediction errors) and that doing so may lead to unexpected interactions between how the environment is represented and what policy is learnt.

The recent success of deep networks trained end-to-end with TD-learning at playing various games has provided a demonstration of how effective this form of learning can be (Mnih et al., 2015; Silver et al., 2016). Moreover, previous work has shown that dopamine neurons project widely to a large number of neural circuits in the brain and end to end learning has been shown to reproduce neural activity in PFC in a wide range of tasks (Song et al., 2017; Wang et al., 2018). This success underscores the importance of understanding how representations learnt directly to maximise overall rewards (or minimise reward prediction errors) may be different than those obtained by maximising other quantities such as reconstruction error. In the context of a rigorously controlled task, our work shows how representations that are inaccurate in encoding several aspects of the task but allow the agent to preserve overall rewards obtained while being representationally efficient can lead to behaviour and reward expectations that are qualitatively different than those that would result from using representations that may best summarise the statistics of the environment or features of the optimal value function and policy.

In sum, by investigating behaviour and DA activity during a time-based decision making task using RL, we were able to reveal an efficient strategy animals appear to be using to represent task variables. We demonstrate that constraints of representational efficiency affect the nature of reward expectations learnt during this task and that the activity of dopaminergic neurons could only be explained by the model using this efficient representation. Finally, we show how animals’ behavioural strategy interacts with the representation used to encode the task in an unexpected way and that this interaction was central for the RL agent to be able to reproduce animals’ behaviour and DA activity. These findings provide novel insights into the manner in which efficiency constraints might be expressed in the reward system, and more generally provide insights into the principles underlying natural, intelligent behavior.

## Acknowledgements

We thank Joao Semedo and Alfonso Renart for several helpful discussions and feedback on the manuscript. We also thank Severin Berger for feedback on the manuscript. This work was developed with the support from the research infrastructure Congento, co-fnanced by Lisboa Regional Operational Programme (Lisboa2020), under the PORTUGAL 2020 Partnership Agreement, through the European Regional Development Fund (ERDF) and Fundação para a Ciencia e Tecnologia (Portugal) under the project L SBOA-01-0145-FEDER-022170. The work was funded by an HHM International Research Scholar Award to J.J.P (#55008745)., European Research Council Consolidator grant (#DYCOC RC - REP-772339-1) to J.J.P., a Bial bursary for scientific research to J.J.P. (#193/2016), internal support from the Champalimaud Foundation. The work was also supported by N H U01 (#NS094288) to C.K.M. and Fundacao para Ciencia e Tecnologia (#SFRH/BD/52214/2013) to A.M.

## Author contributions

AM, JP and CM designed the study. AM performed all analyses and simulations. SS, BA and JP designed and performed the behavioral and photometry experiments. AM, JP and CM wrote the manuscript. JP and CM contributed equally to this work.

---

**Supplementary figure S1:**
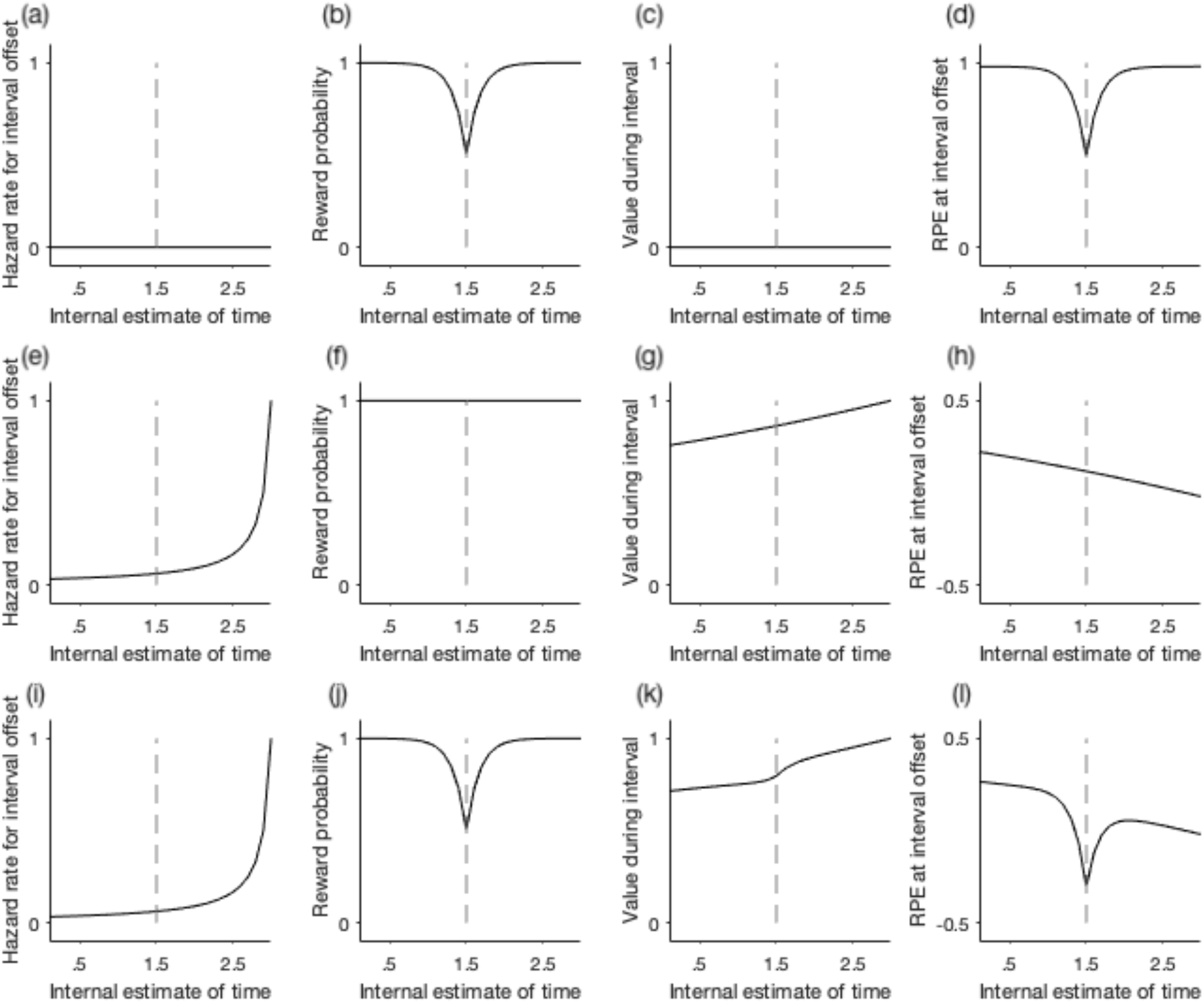
Schematic to illustrate how various task events influence reward expectations. Let’s assume the duration of intervals presented is uniformly distributed between O to 3 sec. and that choices are reported only at interval offset. In this case, the agent needs to estimate value functions over two sets of states, one during the interval and the other at interval offset. Let’s assume the reward amount is 1, in this case the value at interval offset C(z) will be equal to the probability of reporting the correct choice for that interval estimate. During the interval, the value function is a weighted sum of the value of interval offset at that time and the value of being in the interval at later times. The weighting factor is given by the probability of transitioning to each of these states, i.e. V(z) = p(z’ = interval offset | z).C(z) + y.p(z’ ≠ interval offset | z)).V(z’), where z’ is the successor state. **(a-d)** If we assume that the agent does not explicitly encode the distribution of interval durations, but does encode an estimate of choice accuracy at interval offset, the hazard rate encoded by the agent is zero for all internal estimates of time (shown in panel a), and its estimate of value during the interval will always be zero (shown in panel c). Its estimate of value at interval offset is equal to the probability with which it will correctly report interval duration (shown in panel b) and the resulting RPEs at interval offset will reflect only choice accuracy (shown in panel d). **(e-h)** Let’s now assume that the agent encodes the distribution of interval durations, but does not encode choice accuracy. The probability of detecting interval offset at any estimated time z given that no interval offset was detected for all z’ < z is given by the hazard rate of interval offset, H(z). In the case of uniformly distributed interval durations, H(z) is shown in panel e. In this case the value function during the interval is monotonically increasing. If there was no time discounting (i.e. y = 1), this value function would be 1 for the entire interval. For time discounted rewards (i.e. 0 ≪ y < 1), the value function simply reflects the fact that early in the interval the agent expects rewards to be, on average, further in the future and hence more time-discounted than later in the interval (as shown in panel g). Since we assumed here that the agent does not encode choice accuracy, reward expectations from interval offset states are constant (panel f). Consequently, RPEs at interval offset will be monotonically decreasing with elapsed time (as shown in panel h). **(i-1)** Finally, if we assume that the agent encodes, both, choice accuracy as well as the distribution of interval durations, we see that the estimated value at interval offset is the same as when the agent only encodes choice accuracy (shown in panel j). However, the value function during the interval now reflects a combination of choice accuracy and the hazard rate of interval offsets, i.e. V(z) = H(z).C(z) + y.(1-H(z)).V(z’) (shown in panel k). Consequently, the RPEs will also reflect, both, choice accuracy as well as the hazard rate of interval offset (as shown in panel l).

**Supplementary figure S2:**
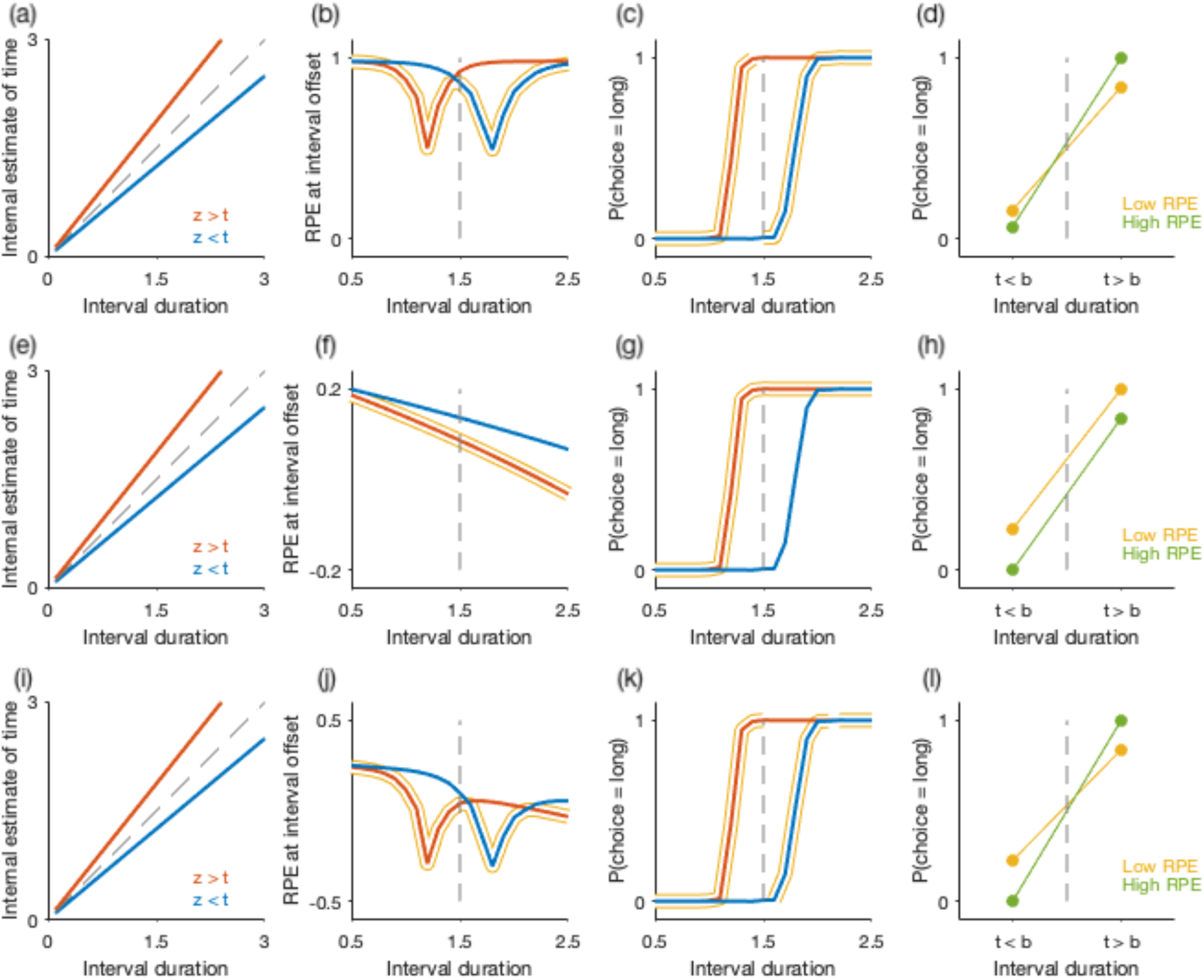
Schematic to illustrate how the overall profile of RPEs at interval offset determines differences between psychometric functions for low and high RPEs trials. Each row shows how over or underestimating elapsed time (shown in column 1) changes RPEs at any possible interval duration (column 2), the probability of reporting choice ‘long’ (column 3) and how magnitude of RPEs relate to the probability of choice ‘long’ (column 4). The top row shows the case when reward expectations are estimated only based on choice accuracy (as shown in Supp Fig 1a-d), the middle row shows the case when reward expectations only reflect the hazard rate of interval offset (as shown in Supp Fig 1e-h) and the bottom row shows the case when reward expectations are estimated based on choice accuracy as well as hazard rate of interval offset (as shown in Supp Fig 1i-l). **(Column 1)** Let’s consider two example types of trials (shown in panels a,e,i), one in which elapsed time is underestimated (z < t, shown by the blue line) and the other overestimates elapsed time (z > t, shown by the red line). **(Column 2)** For these two types of trials, RPEs at interval offsets for all possible interval durations are shown in the second column (each row corresponds to each of the three possibilities for how the agent might estimate reward expectations as a function of task events shown in Supp Fig 1d,h,l). In each panel, for every time step, the trial on which RPE is lower than the other is highlighted in yellow. In panel **(b)**, when RPEs result from reward expectations that only take into account choice accuracy, for all time points before the decision boundary, trials on which elapsed time is overestimated will have lower RPEs than trials on which elapsed time is underestimated. On the other hand, for all time points after the decision boundary, trials on which elapsed time is underestimated will have lower RPEs. In panel **(f)**, when RPEs are driven only due to the hazard rate of interval offset, for all time points trials on which elapsed time is overestimated will have lower RPEs. Finally, in panel **(j)**, when RPEs reflect both choice accuracy and hazard rate of interval offset, RPEs are lower on trials on which elapsed time is overestimated on the short side of the boundary. On the long side of the boundary, close to the boundary, trials on which elapsed time is underestimated have lower RPEs. However, for estimates much longer than the boundary, we see trials on which elapsed time is overestimated have lower RPEs. **(Column 3)** If the agent’s choices change as a function of it’s internal estimates of elapsed time, for the two example trial types shown here, the psychometric function of the agent will also be different. When the agent underestimates elapsed time (blue curve), the psychometric curve will be biased towards ‘short’ choices (i.e. it will show a rightward shift). Similarly, if the agent overestimates time (red curve), the psychometric curve will be biased towards ‘long’ choices (i.e. will be shifted left). To establish the relationship between magnitude of RPE at any given estimated time of interval offset and the probability of choices the agent will report, for each of the trial types, all time points at which RPE was lower on that trial type (shown by the yellow highlights in the second column) are also highlighted in yellow. In panel **(c)**, we see that for all time points before the boundary, the probability of choosing long is higher for most segments highlighted yellow. For all time points after the decision boundary, we see that the probability of choosing short is higher for all segments highlighted in yellow. In panel **(g)**, we see that for all interval durations, trials that had lower RPEs have a higher probability of reporting the interval as ‘long’. Finally, in panel **(k)**, we see that for all interval durations before the boundary, low RPE trials have a higher probability of reporting choice ‘long’ and the opposite is true after the decision boundary. **(Column 4)** For each of the panels c,g and k, for all time points on either side of the boundary we ask: what is the average of the psychometric curves highlighted in yellow. **(d)** In panel c, we see that on the short side of the boundary, for most time points the red psychometric curve is highlighted and the average of the highlighted segments of the curve is shown by the solid yellow marker in the top panel in column d. For time points longer than the boundary, we see in panel c that for most time points, the blue curve is highlighted and the average of that segment of the psychometric curve is shown by the yellow marker on the long side if the boundary in panel d. The green points in panel d show the averages of the curves in panel c on either side of the boundary that are not highlighted and correspond to time points at which the RPEs (shown in panel b) are higher. In other words, when RPEs are driven only due to choice accuracy, lower RPE in general are associated with low choice variability and hence, we would predict the psychometric curves for low and high RPE trials, in this case, to show a difference in slope. **(h)** Following the same steps for the middile row (pane s e-h), we find that when RPEs are driven only by the hazard rate of interval offset, low RPEs correlate with higher probability of reporting long irrespective of which side of the boundary the interval offset lies. Thus, in this case we would predict a change in bias in Psychometric functions for low and high RPE trials. **(I)** Finally, when RPEs reflect both choice accuracy as we as the hazard rate of interval offset, following the same steps we find that the Psychometric curves for low and high RPE trials show a change in slope.

**Supplementary figure S3:**
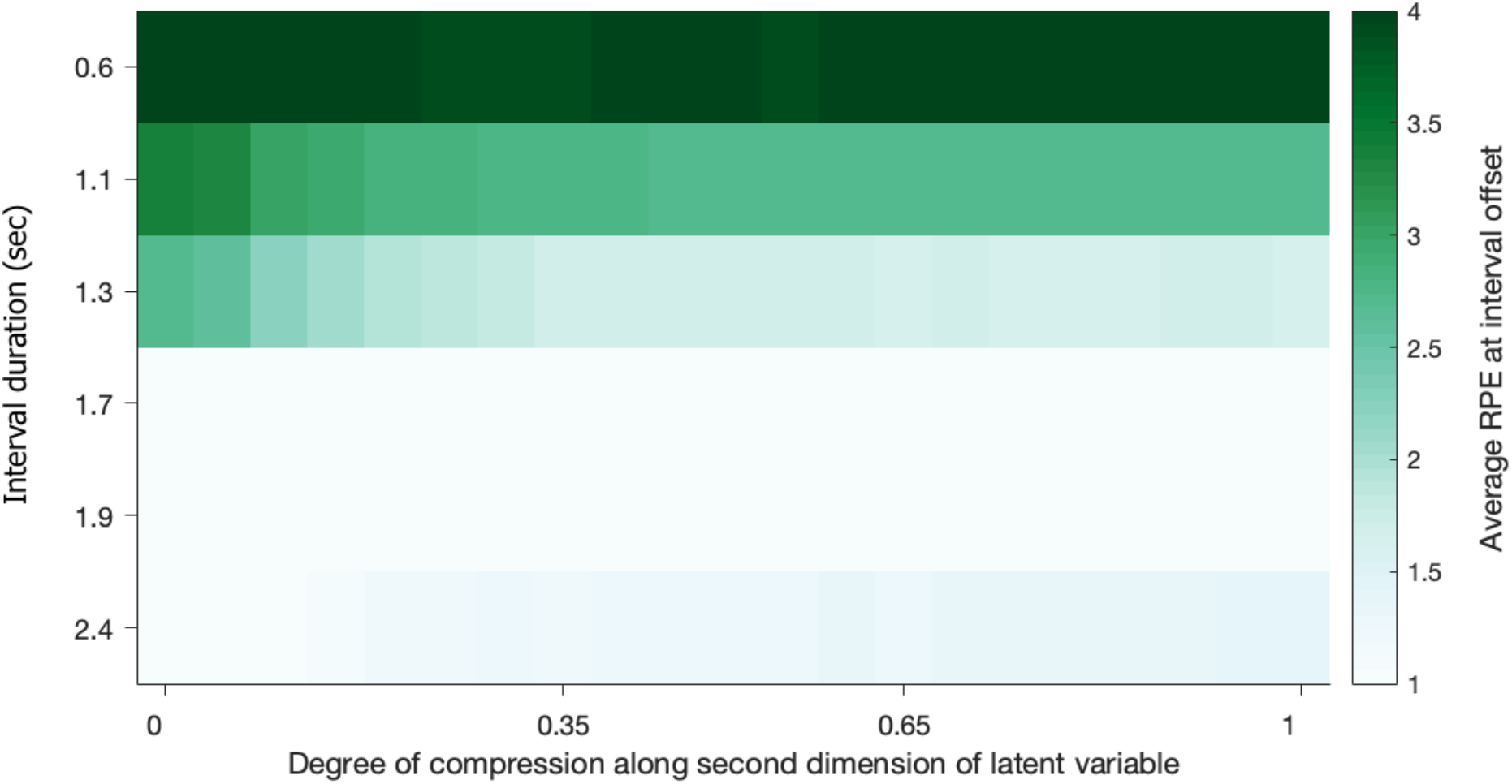
Average reward prediction errors at interval offset for varying degrees of compression in mapping. We see that the profile of average RPEs does not vary considerably for the different degrees of compression in the basic functions used to estimate value functions.

**Supplementary figure S4:**
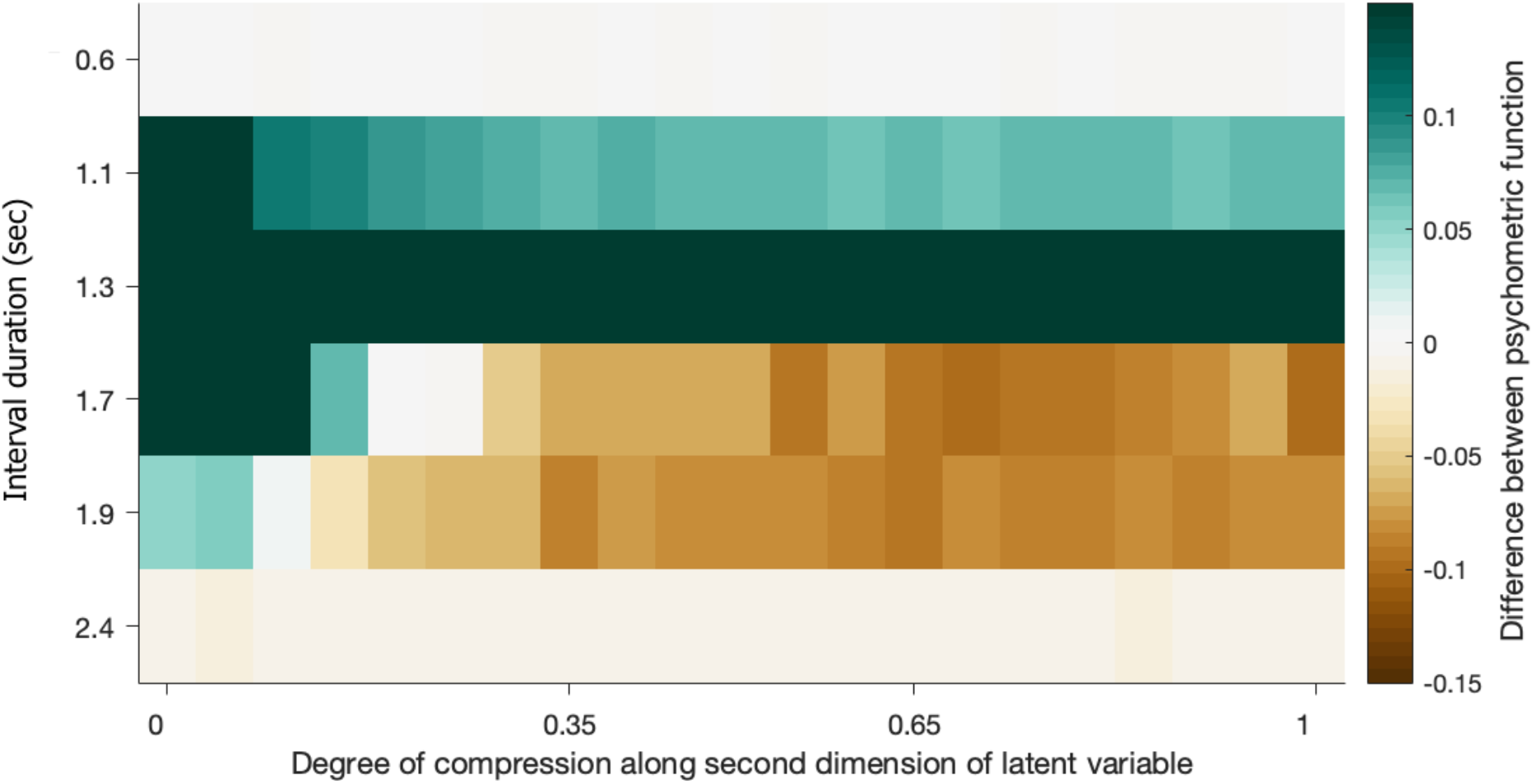
Difference in psychometric functions of trials grouped based on the magnitude of RPEs, for varying degrees of compression in mapping. We see that for the most compressed mapping (DoC = 0), the difference in the psychometric curve has the same sign of all the stimuli presented. This corresponds to a change in bias between the two psychometric functions. On the other hand, for the full mapping (DoC = 1), the difference in the psychometric function changes sign for stimuli on different sides of the boundary. This corresponds to a change in slope between the two psychometric functions. We see that only for mappings that are very close to the most compressed case do the psychometric functions show a difference in bias as is observed in the data.

**Supplementary figure S5:**
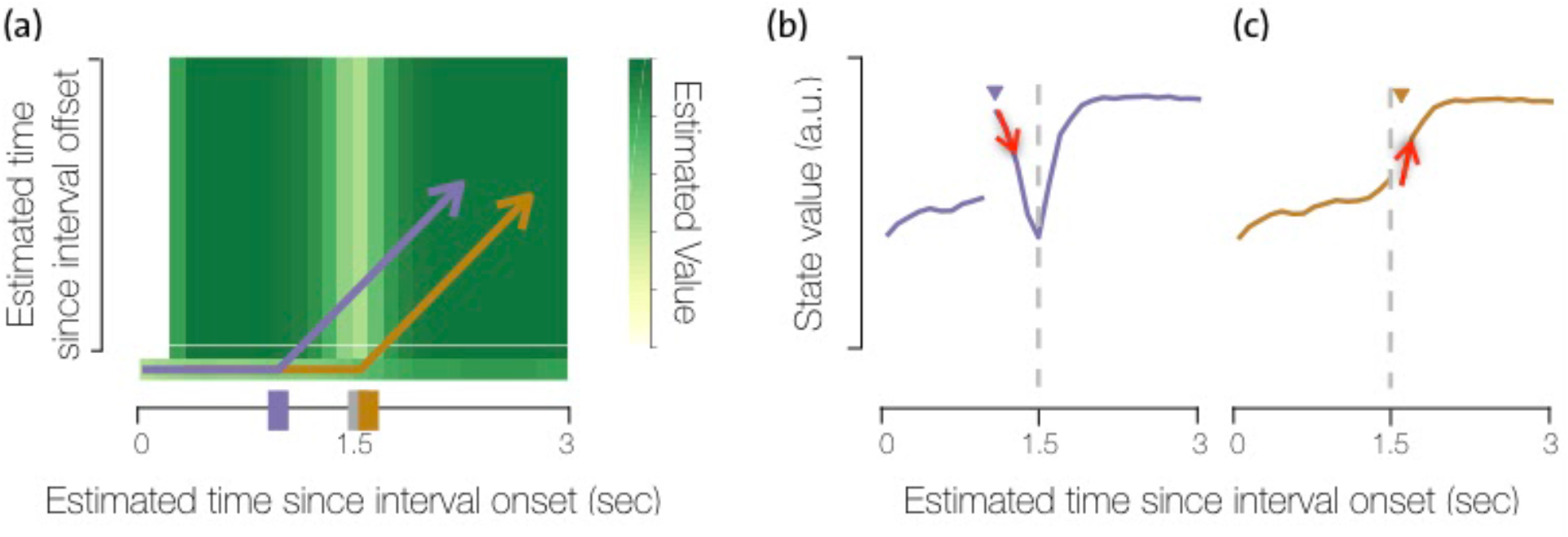
The efficient mapping incentivises procrastination of choices for long interval estimates close to the decision boundary. **(a)** The heat map shows the value function learnt using the efficient mapping when the agent is required to report choices immediately after interval offset. The purple and yellow lines show example trajectories the agent would take through the state space if it withheld choice for the entire trial for an example ‘short’ (purple) and ‘long’ (yellow) interval. **(b-c)** The sequence of state values that the agent will encounter when it follows the purple and yellow trajectories, respectively, shown in (a). The triangle markers indicate the timesteps at which the agent encountered interval offset during these trials. Let’s denote the interval offset states as z_1_ and the subsequent state that the agent transitions to as z_2_. The red arrows show the transition between z_1_ and z_2_ for the example trajectories shown in (b-c). **(b)** We see that after an estimated ‘short’ interval, withholding choice would result in the value function to decrease and hence incur negative RPEs. However, the average reward obtained from reporting a choice immediately would be equal to the value of the interval offset state z_1_ and will, on average, incur zero RPEs. Hence the agent is discouraged from withholding choice when estimating an interval to be ‘short’. **(c)** On the other hand, for intervals estimated as near-boundary ‘long’, withholding choices results in incurring positive RPEs. If the agent reported a choice immediately, it would on average receive rewards equal to the value function at z_1_ and incur zero RPEs in doing so. Hence the agent is incentivised to withhold choices for long near-boundary interval estimates. On these trials, when the agent transitions from z_1_ to z_2_ by withholding choice actions, the value of z_1_ will be updated closer to the value of state z_2_ based on the TD update equation V(z_1_) ← V(z_1_) + *a*(γV(z_2_) − V(z_1_)). Moreover, when the agent reports a choice action at z_2_ after transitioning from z_1_ to z_2_, the average reward it will receive from z_2_ will be lower than if the interval offset had been presented at z_2_. Trial to trial variability in the latent variable z will lead to incorrect estimates of the category (‘short’ vs ‘long’) of intervals presented closer to the decision boundary (such as z_1_) than those further away from it (such as z_2_). Consequently, trial to trial variability in choices at z_2_ will be lower if those choices are from trials where the agent’s estimate at interval offset is z_2_ than if choices at z_2_ are reported on trials on which agent’s estimate at interval offset is either z_2_ or z_1_. Thus, TD-updates on the long side of the decision boundary lead to a flattening of the value function in the efficient model due to procrastination of choices.

**Supplementary figure S6:**
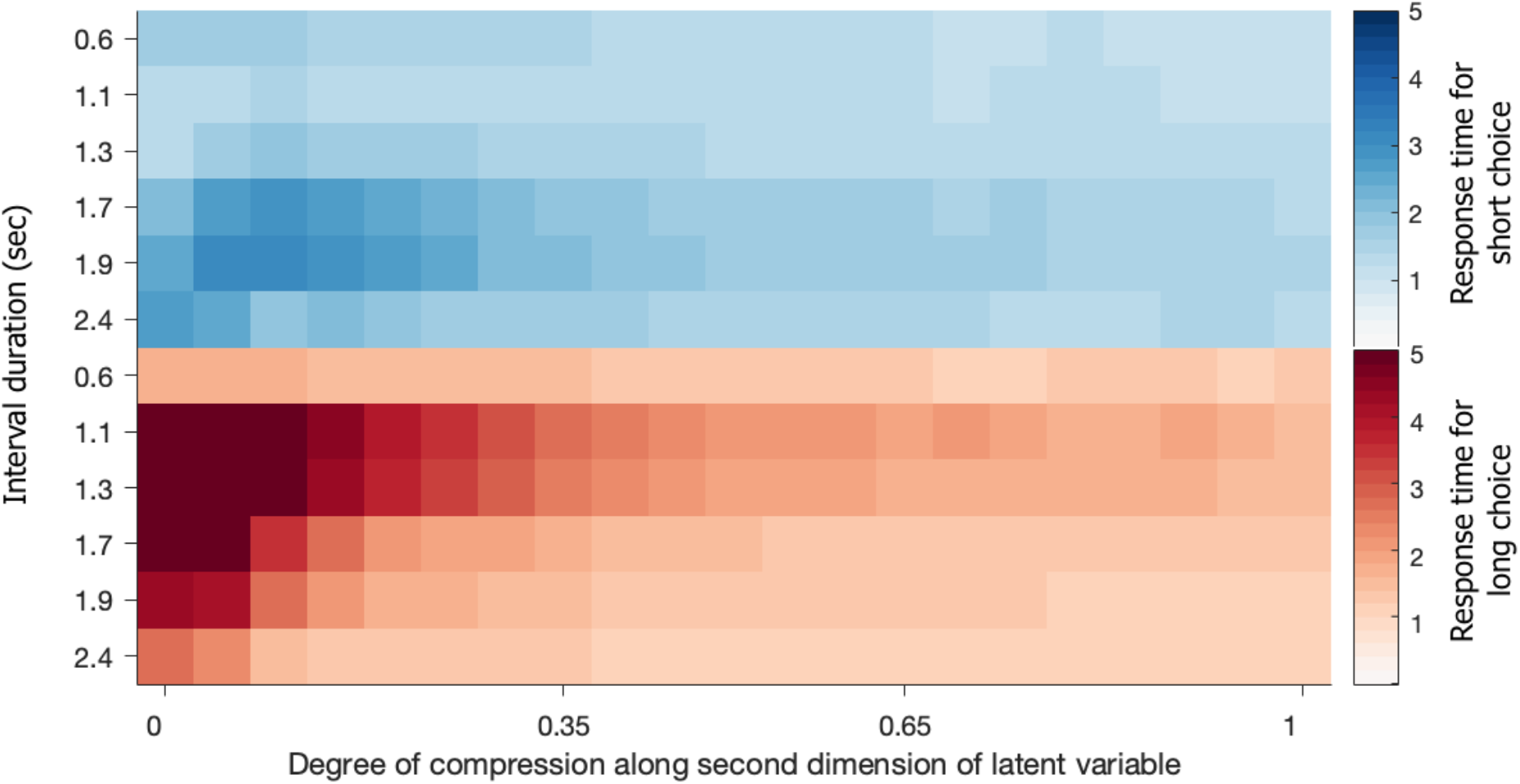
Average response times grouped based on the agent’s choice for varying degrees of compression in mapping. The agent has short response times for all stimuli when choosing short. The profile is very similar for all degrees of compression in the mapping used. However, the profile of average response times when the agent chooses long changes greatly with the degree of compression in the mapping. Only when using very compressed mappings do we see response times decrease with interval duration on trials with ‘long’ choices as seen in the data.

## Methods

Here we describe how we modelled reinforcement learning (RL) agents on the interval discrimination task and obtained reward prediction error signals that we compared against observed dopamine activity recorded in animals behaving in a similar task. We will organise the following section into: (1) how we define the state representation for the agent and what is the set of permitted actions the agent can choose from; (2) how, given the state representation, the agent estimates reward expectations and an appropriate mapping between states and actions; (3) parameters with which we simulated our RL agents; (4) an equivalent way in which the compression of the efficient representation could be implemented.

### 1 State representation encodes elapsed time since observed task events

During the interval discrimination task (IDT), animals and agents are provided with a very sparse set of cues or observations. Interval onset and offset are both signalled by identical tones, and animals have to infer from context the start, end and duration of intervals presented to them. Animals also observe whether or not their choices result in rewards or no rewards, based on which they need to learn to make appropriate choices. Hence, the RL agent needs to have an internal representation that allows it to use the above described observations to infer the correct epoch in the task (i.e. to infer at any given time if the agent is in the stimulus interval, after the interval or in the inter-trial-interval) as well as estimate interval duration presented. The interval duration presented on each trial is randomly sampled uniformly from a set of six intervals and this re the stimulus distribution used in task for the animals. Finally we model all times during inter-trial intervals with a single terminal state i.e. a state that always has zero reward expectations. In other words, we model the task as episodes and each trial is a new episode for the agent.

Animals trained on the task are not restrained during the interval and any report of choice during the interval results in a premature termination of the trial, accompanied with no reward and a time-out to keep the trial-rate constant. Hence animals need learn to not only classify the presented interval with the appropriate choice, but they need to learn to withhold reporting any choice until after the end of the interval. Similar to animals, the RL agent is permitted to make one of three actions at any time during the task: report choice *short*, report choice *long* and withhold both choices i.e. *wait*. Like animals, the agent needs to learn not only report which choice to make, but when to make it. The agent need to learn to withhold choices during the interval and report a *short* or *long* choice based on its estimate of interval duration, after interval offset. We model the task such that episodes always begin at interval onset and terminate when the agent reports either of the two choices or until the trial time-out.

Specifying a RL agent that encodes reward expectations as a function of internal representations that evolve both based on internal variables as well as observations from the environment (i.e. task) can be formally described as partially-observed Markov decision process (POMDP). In a POMDP, at any time, the agent combines a likelihood based on the observations it gets from the environment with a prior that it computes from earlier estimates of its internal variables. This results in a posterior estimate or belief over the agent’s internal variable using which the agent can then estimate reward expectations. Since POMDPs provide a probabilistic framework with which an agent can combine internal variables and observations from the environment, its resulting beliefs are also in the form of probability distributions. Exactly computing reward expectations as based on distributions of internal states is, in general, an extremely challenging problem. There are several algorithms that use different types of approximations to solve POMDPs, but it is unclear how computations on POMDPs might be implemented in neural circuits. Our model can be described as one in which the agent does not maintain an explicit distribution over its internal variables, but one in which its internal variables evolve such that the agent’s states are samples from the posterior distribution or belief over these states. These internal variables are often referred to as latent variables or states.

#### 1.1 Dynamics of the latent variable

The task requires the latent variable dynamics to have two key features: (1) when receiving no observations from the environment, the latent variable needs to evolve, as time elapses, in a non-repeating manner, such that the latent variable can provide an internal estimate of elapsed time; (2) the latent variable must change as a function of interval onset and offset, as well as agent’s actions, such that it reflects the task epoch at any point. Accordingly, the agent encodes a continuous two-dimensional latent variable **z** ∈ ℝ^2^, where each dimension represents an estimate of elapsed time since interval onset and offset, respectively.

Additionally, we know that animals’ estimates of elapsed time vary from trial-to-trial, and that variability in their choices is driven by variability in how far their neural activity evolves during the interval [4] Hence, we modelled the evolution of the latent variable to also have trial-to-trial variability in its dynamics. There is a large class of stochastic dynamical systems that can fulfil these requirements. One simple way of fulfilling (1) and (2), and introducing variability into the state representation is to de a linearly accumulating two-dimensional stochastic system. When the agent withholds choices (i.e., when the agent *waits*), the evolution of the latent variable is, thus, governed by two factors: (1) Observations in the task: accumulation along the first dimension only begins after interval onset and during the interval the latent variable along the second dimension remains zero. At interval offset, accumulation begins along the second dimension as well; (2) Internal noise: the accumulation along each of the dimensions after interval onset and offset, respectively, is noisy and has trial to trial variability.

More specifically episodes always begin at interval onset, and the initial state is given by **z**_0_ = [0, 0]^**T**^. Whenever the agent makes a *choice* (*left* or *right*), the latent variable transitions to the terminal state **z**_**T**_ = [∞, ∞]^**T**^, which marks the end of the current episode. During the interval, dynamics are de as:

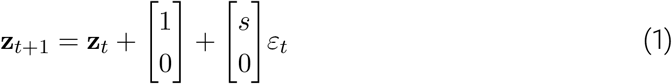

where 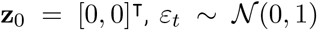, and *s* controls the standard deviation of the additive noise. Previous work has shown that trial to trial variability in estimates of elapsed time also follow Weber’s law and in the time domain, this is referred to as the scalar property, i.e. the standard deviation of estimates of elapsed time increase linearly with elapsed time [3] In order for the latent variable encoded by the RL agent to exhibit scalar variability, we specify the standard deviation of additive noise to be 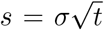, where σ = .25. Thus, the standard deviation of the latent variable along the first dimension 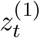, at time *t* during the interval, will be σ*t*. The value of parameter σ was chosen to allow the psychometric function of choices reported by the RL agent to qualitatively match those of animals trained on this task.

At interval offset (when the tone occurs), we increment the latent variable along the second dimension with an increment of fixed size to allow the observation of interval offset to be reliable encoded by the latent variable. Thus, we have:

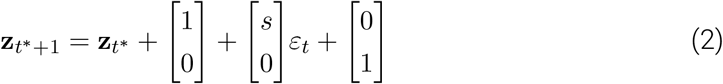

where *t*\* indicates the time step at which interval offset occurred. After which, the latent variable evolve as follows:

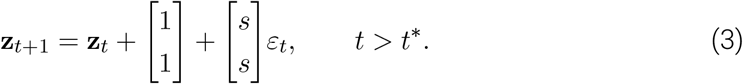

### 2 Learning

In the simulated task, the agent receives a large positive reward for correctly reporting the interval category after interval offset. Incorrect choices after interval offset and any choice during the interval incur a small negative reward. The goal of an agent is to estimate, for each state, (i) a state-value function, which is an estimate of expected discounted future rewards from that state, and (ii) a policy, which is a probabilistic mapping between state and actions that maximise overall future rewards. We model the agent to learn value functions using TD-learning within an actor-critic architecture since they have been commonly used to model dopamine activity as RPEs. The state-value function is referred to as the *critic* since it evaluates the value of each state. And the *actor* encodes the policy with which the agent makes actions during the task.

#### 2.1 State value estimation

When the state of the environment can be observed and is discrete, the expectation of discounted future rewards, referred to as *value*, is de for each state the agent can encounter in its environment, and can be written recursively in terms of value of future states [9].

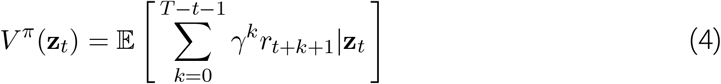

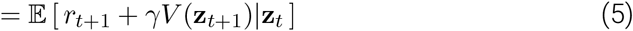

where the expectation is with respect to, both, the dynamics of the environment and the agent’s policy π, which the agent samples as it interacts with the environment. However, since the latent variable in our RL agent is continuous, we cannot innumerate a value for each possible instance of the latent variable and require an approximation scheme to estimate value. Hence, we map the latent variable at any given time **z**_*t*_ into a D-dimensional feature space, such that linear combinations of the features can be used to approximate the value function for all instances of **z**_*t*_.

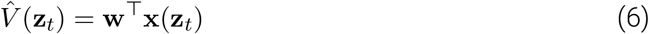

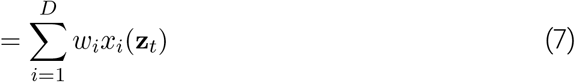

where the components of the feature vectors or basis functions **x** = [*x*_1_, …, *x*_*D*_] are constructed using tile basis. The parameters of the basis functions *θ*_*i*_ = {*b*_*i,l*_, *b*_*i,u*_, *c*_*i,l*_, *c*_*i,u*_} determine the range of **z** over which each component of the bases *x*_*i*_ is non-zero, as shown below:

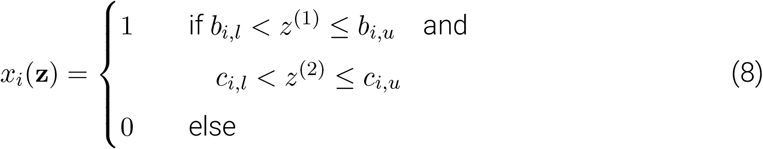

To estimate **w**, that best approximates the state value function, we make updates to reduce the loss function given by

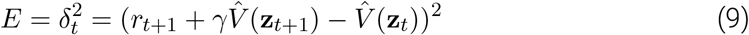

where *δ*_*t*_ is referred to as the temporal-difference (TD) error. The updates are proportional to the gradient of this loss function, where we treat the target 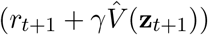 as a constant with respect to the parameters *w*_*i*_, as shown below:

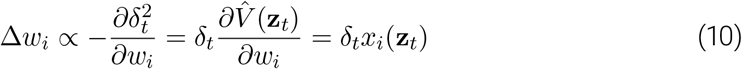

Hence, we can write the update rule for each component of the weights vector **w** as

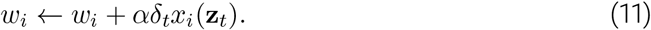

#### 2.2 Policy learning

The ultimate goal of a RL agent is to learn a policy that maximises its future expected rewards. To allow our agent to appropriate learn state-actions mappings, we model the actor to estimate the advantage function for every action *A*(**z**, *a*) [1].

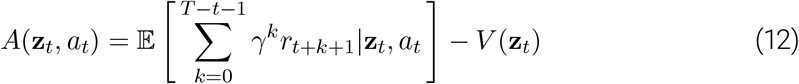

where, the first term i.e. the expected cumulative future rewards for a given state-action pair is referred to as the state-action value function *Q*(**z**, *a*). Hence, the advantage function, *A*(**z**, *a*) = *Q*(**z**, *a*) − *V* (**z**), quantifies the difference or advantage in expected returns of taking a particular action in a given state as compared to the overall expected returns from that state. For a given state-action pair, the advantage function can be estimated using updates based on the TD-error computed by the critic (Equation 9) as shown below:

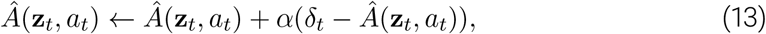

As in the case of the state value function, we approximate 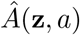 as a linear combination of feature vectors and we update the parameters of this function using the TD-error as described below:

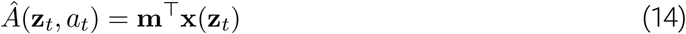

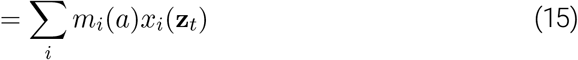

The update rule to estimate parameters **m**, is similarly given by

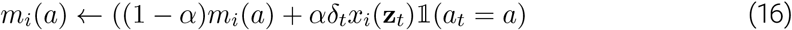

Finally, at every step, the actor samples an action from the policy distribution π(**z**, *a*)= *p*(*a*|**z**), which is given by a soft-max of function of the approximation of the advantage function 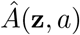, as shown below:

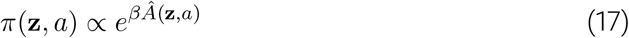

### 3 Model parameters

#### 3.1 Parameters of basis functions for value estimation

The tile bases used in the model implementations are fully described by the range over which they are 1. We keep the width of the basis function along *z*^(1)^ the same for all versions of the model and we only vary the width of the basis function along *z*^(2)^. The width of basis function along *z*^(1)^ was chosen to allow an accurate approximation of value along this axis given the trial to trial variability in estimates of elapsed time. Since the tile bases are non-overlapping, we can describe the parameters of all the basis functions in terms of the boundaries along the two dimensions of **z**.

Let the boundaries along the first dimension of the latent variable, which encodes elapsed time since interval onset, *z*^(1)^ be *B* = {0, *b*_0_, *b*_1_, …*b*_*K*_}. The resolution of the value function approximation the same along the first dimension is not varies for different degrees of compression in the mappings. The tiling along this dimension is also kept uniform and the interval between the boundaries is set to be equal to ms. Let the boundaries along the second dimension of the latent variable, which encodes elapsed time since interval offset, *z*^(2)^ be *C* = {0, *c*_0_, *c*_1_, …*c*_*K*_}. In the full mapping, the parameters of the tile bases along *z*^(1)^ and *z*^(2)^ are the same. The compression in the mapping only in the value function along this second axis and hence the set of boundaries in *C*. Let the compression factor be λ [0, 1], such that the full mapping corresponds to λ = 1 and the efficient mapping (the most compressed representation) corresponds to λ = 0. In this case, *c*_*i*_ = *b*_*i*_/(1 − λ), ∀ *i* ∈ {1, 2, …*K*}.

#### 3.2 Other parameters

The value function estimated by the agent is de as the estimate of future discounted rewards from any state and we set the temporal discounting parameter γ = .95.

Since the parameters for estimating the state value function as well as the advantage function are learnt incrementally from the agent’s interaction with the environment, we needed to specify the learning rate for each of these sets of parameters. We implemented an actor-critic framework and the learning rates were chosen such that the updates of the actor occur on a slower timescale than those of the critic. This is to allow for the critic to have enough updates to evaluate the current policy. We specify the learning rate for the state value function to be *α* = 1/*n*_*i*_ when the number of visits to the state being updated *n*_*i*_ is less than *N*_*V*_ = 100, and *α* = 1/*N*_*V*_ otherwise. The learning rate for the advantage function is similarly set to be *α* = 1/*n*_*i*_ when the number of visits to the state being updated *n*_*i*_ is less than *N*_*A*_ = 1000. Thus, after the first few visits to any state, the learning rate for the state-value function is much faster than that of the advantage function. In the current work, we are interested in relating value-based behaviour and RPEs after the the agent has fully learnt the task to behaviour and neural activity in over-trained animals on the interval discrimination task. Hence, the learning parameters were chosen without any consideration to model the time-course of learning of animals on this task.

After learning has converged, there are two sources of trial-to-trial variability in the agent’s behaviour in our implementation: variability in dynamics of the latent variable and variability due to stochasticity in the policy. The parameter for the standard deviation of additive noise was set to be σ = .25 and the parameter that determines the stochasticity of the policy was chosen to be *α* = 3. Both these parameters were held constant over the entire duration of learning, for all simulations shown.

### 4 Alternative formulation for the ef representation consistent with observed dopaminergic activity and animals’ behaviour

Our use of term ‘representational efficiency in estimating the value functions can be described in one of two equivalent ways. First, representational efficiency corresponds to a reduction in the number of non-zero parameters that need to be estimated to approximate the value function over all possible instances the latent variable can take during a given task. Alternatively, representational efficiency can be described in terms of the number of basis functions that will be non zero while representing all possible instances the latent variable can take for the given task. For any linear function approximation scheme such as 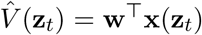, this refers to reducing the dimensionality of **w** and **x**. This can be done by either changing the dynamics of the latent variable, while maintaining the parameters of the basis functions constant, or by changing the parameters of the basis functions while maintaining the dynamics of the latent variable constant.

In the first case, for a fixed set of basis functions, the dynamics of **z** can be changed such that *p*(**z**) is non-zero in a smaller area of state space. In this case fewer basis functions will be needed to estimate the value function over all possible instances of **z**_*t*_. Alternatively, if the dynamics of the latent variable are kept constant, representational efficiency can be achieved by changing the range over which each basis function is non-zero, such that individual basis functions span larger regions of the state space i.e. the space of **z**. Moreover, both, the dynamics of the latent variable and well as the parameters of the basis function can be changed simultaneously to achieve compression.

The RL agent with the efficient mapping presented in this work can similarly be refor-mulated by specifying modified dynamics of the latent variable as opposed to by the parameters of the basis functions (as described in the main text), or by changing both. The equivalent formulation can be expressed by introducing the compression factor λ ∈ [0, 1] in the dynamics of the second latent dimension as follows:

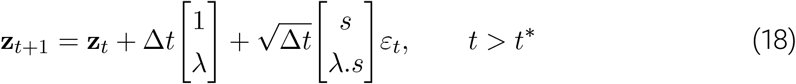

The full or uncompressed representation corresponds to λ = 1, while the efficient representation (where the second latent variable does not perform any accumulation) corresponds to λ = 0. Note that changing λ does not have any effect on the state representation during the interval. Rather, after the interval offset, as λ → 0 the state representation loses the ability to resolve the time of interval offset and is only able to encode time since interval onset and whether or not interval offset has been presented or not.

In either formulation, as long as the relationship between the latent variable and the basis functions is maintained, our results are invariant to which of the two sets of variables (the dynamics of the latent variable *p*(**z**) or the basis functions for value approximation **x**) are changed to achieve a reduction in parameters to be estimated. Hence, for simplicity, in the main text we focus on describing our results for the formulation in which the dynamics of the latent variable are held constant and the density of the basis functions along the second dimension is reduced.

### 5 Alternative ways of obtaining ef representations that do not reproduce dopaminergic activity and animals’ behaviour

Similarly, there are multiple ways in which the dynamics of the latent variable and the basis functions can be changed to yield a function approximation that has equivalent number of parameters as the one discussed above, but would not have the same relationship between the latent variable and the basis functions discussed. However, none of these alternatives were able to simultaneously reproduce the average pro of dopamine activity and the trial to trial relationship between magnitude of dopamine response and temporal judgements. These alternatives include:

1. Fast reaction times for all estimates of interval offsets: If the policy of the agent is always report choice immediately after interval offset, the latent variable would only span the first couple of basis functions along *z*^(2)^ and hence would require similar number of parameters as the efficient mapping. In this case, both the latent variable and the bases for value approximation remain the same. However, by only allowing the agent to choose between actions associated with ‘short’ and ‘long’ choice at interval offset, the latent variable does not … The reward expectations obtained in this version are the same as when we directly computed reward expectations based on choice accuracy and predictability of interval offset in time (shown in Fig 2n and Supp. Fig. 2i-j). However it is known that animals will forgo some amount and immediacy of rewards to balance physical effort in obtaining rewards
2. Latent variable dynamics such that there is no change in the state space after interval offset: If the latent variable encoded by the agent does not change with time after interval offset, it can serve as a perfect ‘memory’ of the estimated time of interval offset. In this case, even though the function approximation is the same as that used in the full model, only a small number of bases are needed to encode all possible instances of the latent variable. The reward expectations learnt using this version for all states during the interval and at interval offset are qualitatively the same as those learnt using the full model. This alternative requires the latent variable dynamics to change very abruptly from sequential to attractor dynamics at interval offset. We can speculate that such an abrupt change in dynamics of the latent variable may be hard to achieve if we assume some smoothness constraints in the dynamics neural circuits can generate [8].
3. Latent variable dynamics such that there is no change in the projection of the latent variable on the axis that encodes elapsed time since interval onset after interval offset: Previous work has shown that neural circuits can indeed be con such that irrelevant activity falls in the null space of the required readout [2,5,6]. This version would leave the function approximation the same as in (1) and would produce the same results and the time evolution of the latent variable during the interval would also remain the same. The key difference here would be in the dynamics of the latent variable after interval offset and can be described as follows:

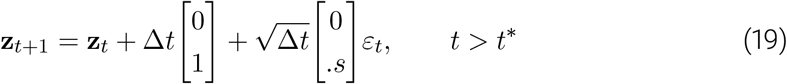

This formulation allow an unambiguous approximation of the state of the task. In the current task, the observations available to the animal may be too sparse to allow neural circuits to ne tune dynamics after interval offset.
4. Mapping in which the post interval basis functions are oriented to be parallel to the directions along which the latent variable evolves after interval offset: This alternative would also require that the basis functions for the mapping are finely tuned based on the dynamics of the latent variable.

### 6 Resource availability

Further information and requests for resources should be directed to and will be fullfilled by the Lead Contact, Asma Motiwala.

## References

Bayer, H. M., & Glimcher, P. W. (2005). Midbrain Dopamine Neurons Encode a Quantitative Reward Prediction Error Signal. Neuron, 47(1), 129–141. https://doi.org/10.1016/j.neuron.2005.05.020

Botvinick, M. M. (2008). Hierarchica models of behavior and prefronta function. Trends in Cognitive Sciences, 72(5), 201–208. https://doi.org/10.1016/j.tics.2008.02.009

Botvinick, M. M., Niv, Y., & Barto, A. G. (2009). Hierarchically organized behavior and its neura foundations: A reinforcement learning perspective. Cognition, 113(3), 262–280. https://doi.org/10.1016/j.cognition.2008.08.011

Botvinick, M., Weinstein, A., Solway, A., & Barto, A. (2015). Reinforcement learning, efficient coding, and the statistics of natural tasks. Current Opinion in Behavioral Sciences, 5, 71–77. https://doi.org/10.1016/j.cobeha.2015.08.009

Daw, N. D., Courville, A. C., & Touretzky, D. S. (2006). Representation and Timing in Theories of the Dopamine System. Neural Computation, 18(7), 1637–1677. https://doi.org/10.1162/neco.2006.18.7.1637

Dayan, P., & Sejnowski, T. J. (1994). TD(λ) converges with probability 1. Machine Learning, 74(3), 295–301. https://doi.org/10.1007/BF00993978

Druckmann, S., & Chklovskii, D. B. (2012). Neuronal Circuits Underlying Persistent Representations Despite Time Varying Activity. Current Biology, 22(22), 2095–2103. https://doi.org/10.1016/j.cub.2012.08.058

Fiorillo, C. D., Newsome, W. T., & Schultz, W. (2008). The temporal precision of reward prediction in dopamine neurons. Nature Neuroscience, 11(8), 966–973. https://doi.org/10.1038/nn.2159

Fiorillo, C. D., Tobler, P. N., & Schultz, W. (2003). Discrete Coding of Reward probability and Uncertainty by Dopamine Neurons. Science, 299(5614), 1898–1902. https://doi.org/10.1126/science.1077349

Gershman, S. J., Norman, K. A., & Niv, Y. (2015). Discovering latent causes in reinforcement learning. Current Opinion in Behavioral Sciences, 5, 43–50. https://doi.org/10.1016/j.cobeha.2015.07.007

Gibbon, J., & Church, R. M. (1990). Representation of time. Cognition, 37(1), 23–54. https://doi.org/10.1016/0010-0277(90)90017-E

Gouvêa, T. S., Monteiro, T., Motiwala, A., Soares, S., Machens, C., & Paton, J. J. (2015). Striata dynamics exp ain duration judgments. ELife, 4. https://doi.org/10.7554/eLife.11386

Janssen, P., & Shadlen, M. N. (2005). A representation of the hazard rate of elapsed time in macaque area LIP. Nature Neuroscience, 8(2), 234–241. https://doi.org/10.1038/nn1386

Joel, D., Niv, Y., & Ruppin, E. (2002). Actor-critic models of the basal ganglia New anatomica and computationa perspectives. Neural Networks, 15(4), 535–547. https://doi.org/10.1016/S0893-6080(02)00047-3

Kaufman, M. T., Churchland, M. M., Ryu, S. I., & Shenoy, K. V. (2014). Cortica activity in the nu space Permitting preparation without movement. Nature Neuroscience, 77(3), 440–448. https://doi.org/10.1038/nn.3643

Kepecs, A., Uchida, N., Zariwala, H. A., & Mainen, Z. F. (2008). Neura corre ates, computation and behavioura impact of decision confidence. Nature, 455(7210), 227–231. https://doi.org/10.1038/nature07200

Khamassi, M., Lachèze, L., Girard, B., Berthoz, A., & Guillot, A. (2005). Actor-Critic models of Reinforcement Learning in the Basal Ganglia: From Natural to Artificial Rats. Adaptive Behavior, 13(2), 131–148. https://doi.org/10.1177/105971230501300205

Kiani, R., & Shadlen, M. N. (2009). Representation of Confidence Associated with a Decision by Neurons in the Parietal Cortex. Science, 324(5928), 759–764. https://do.org/10.1126/science.1169405

Lak, A., Nomoto, K., Keramati, M., Sakagami, M., & Kepecs, A. (2017). Midbrain Dopamine Neurons Signal Belief in Choice Accuracy during a Perceptual Decision. Current Biology, 27(6), 821–832. https://do.org/10.1016/j.cub.2017.02.026

Ludvig, E. A., Sutton, R. S., & Kehoe, E. J. (2008). Stimulus Representation and the Timing of Reward-Prediction Errors in models of the Dopamine System. Neural Computation, 20(12), 3034–3054. https://do.org/10.1162/neco.2008.11-07-654

Mello, G. B. M., Soares, S., & Paton, J. J. (2015). A Scalable Population Code for Time in the Striatum. Current Biology, 25(9), 1113–1122. https://do.org/10.1016/j.cub.2015.02.036

Mnih, V., Kavukcuoglu, K., Silver, D., Rusu, A. A., Veness, J., Bellemare, M. G., Graves, A., Riedmiller, M., Fidjeland, A. K., Ostrovski, G., Petersen, S., Beattie, C., Sadik, A., Antonoglou, I., King, H., Kumaran, D., Wierstra, D., Legg, S., & Hassabis, D. (2015). Human-level control through deep reinforcement learning. Nature, 518(7540), 529–533. https://do.org/10.1038/nature14236

Niv, Y., & Langdon, A. (2016). Reinforcement learning with Marr. Current Opinion in Behavioral Sciences, 11, 67–73. https://do.org/10.1016/j.cobeha.2016.04.005

Pasquereau, B., & Turner, R. S. (2014). Dopamine neurons encode errors in predicting movement trigger occurrence. Journal of Neurophysiology, 113(4), 1110–1123. https://do.org/10.1152/jn.00401.2014

Remington, E. D., Narain, D., Hosseini, E. A., & Jazayeri, M. (2018). Flexible Sensorimotor Computations through Rapid Reconfiguration of Cortical Dynamics. Neuron, 98(5), 1005–1019.e5. https://do.org/10.1016/j.neuron.2018.05.020

Reynolds, J. N. J., Hyland, B. I., & Wickens, J. R. (2001). A cellular mechanism of reward-related learning. Nature, 413(6851), 67–70. https://do.org/10.1038/35092560

Roitman, J. D., & Shadlen, M. N. (2002). Response of Neurons in the Lateral Intraparietal Area during a Combined Visual Discrimination Reaction Time Task. Journal of Neuroscience, 22(21), 9475–9489. https://do.org/10.1523/JNEUROSC.22-21-09475.2002

Russek, E. M., Momennejad, I., Botvinick, M. M., Gershman, S. J., & Daw, N. D. (2017). Predictive representations can link model-based reinforcement learning to model-free mechanisms. PLOS Computational Biology, 13(9), e1005768. https://do.org/10.1371/journa.pcb.1005768

Salinas, E. (2006). How Behavioral Constraints May Determine Optimal Sensory Representations. PLOS Biology, 4(12), e387 https://do.org/10.1371/journa.pbio.0040387

Schultz, W., Dayan, P., & Montague, P. R. (1997). A Neural Substrate of Prediction and Reward. Science, 275(5306), 1593–1599 https://do.org/10.1126/science.275.5306.1593

Semedo, J. D., Zandvakili, A., Machens, C. K., Yu, B. M., & Kohn, A. (2019). Cortical Areas Interact through a Communication Subspace. Neuron, 102(1), 249–259.e4. https://do.org/10.1016/j.neuron.2019.01.026

Silver, D., Huang, A., Maddison, C. J., Guez, A., Sifre, L., van den Driessche, G., Schrittwieser, J., Antonoglou, I., Panneershelvam, V., Lanctot, M., Dieleman, S., Grewe, D., Nham, J., Kalchbrenner, N., Sutskever, I., Lillicrap, T., Leach, M., Kavukcuoglu, K., Graepel, T., & Hassabis, D. (2016). Mastering the game of Go with deep neural networks and tree search. Nature, 529(7587), 484–489. https://do.org/10.1038/nature16961

Soares, S., Atallah, B. V., & Paton, J. J. (2016). Midbrain Dopamine neurons control judgment of time. Science, 354(6317), 1273–1277. https://do.org/10.1126/science.aah5234

Song, H. F., Yang, G. R., & Wang, X.-J. (2017). Reward-based training of recurrent neural networks for cognitive and value-based tasks. ELife, 6, e21492. https://do.org/10.7554/eLife.21492

Starkweather, C. K., Babayan, B. M., Uchida, N., & Gershman, S. J. (2017). Dopamine reward Prediction errors reflect hidden-state inference across time. Nature Neuroscience, 20(4), 581–589. https://do.org/10.1038/nn.4520

Stauffer, W. R., Lak, A., & Schultz, W. (2014). Dopamine Reward Prediction Error Responses Reflect Marginal Utility. Current Biology, 24(21), 2491–2500. https://do.org/10.1016/j.cub.2014.08.064

Steinberg, E. E., Keiflin, R., Boivin, J. R., Witten, I. B., Deisseroth, K., & Janak, P. H. (2013). A causal link between Prediction errors, dopamine neurons and learning. Nature Neuroscience, 16(7), 966–973. https://do.org/10.1038/nn.3413

Suri, R. E., & Schultz, W. (1998). Learning of sequential movements by neural network model with dopamine-like reinforcement signal. Experimental Brain Research, 121(3), 350–354. https://do.org/10.1007/s002210050467

Sutton, R. S. (1988). Learning to predict by the methods of temporal differences. Machine Learning, 3(1), 9–44. https://do.org/10.1007/BF00115009

Wang, J., Narain, D., Hosseini, E. A., & Jazayeri, M. (2018). Flexible timing by temporal scaling of Cortical responses. Nature Neuroscience, 21(1), 102–110. https://do.org/10.1038/s41593-017-0028-6

Wang, J. X., Kurth-Nelson, Z., Kumaran, D., Tirumala, D., Soyer, H., Leibo, J. Z., Hassabis, D., & Botvinick, M. (2018). Prefrontal cortex as a meta-reinforcement learning system. Nature Neuroscience, 21(6), 860–868. https://do.org/10.1038/s41593-018-0147-8

Watabe-Uchida, M., Eshel, N., & Uchida, N. (2017). Neural Circuitry of Reward Prediction Error. Annual Review of Neuroscience, 40(1), 373–394. https://do.org/10.1146/annurev-neuro-072116-031109

Wimmer, G. E., Daw, N. D., & Shohamy, D. (2012). General zat on of value in reinforcement learning by humans. European Journal of Neuroscience, 35(7), 1092–1104. https://do.org/10.1111/j.1460-9568.2012.08017.x

## References

[1] Leemon C Baird. Advantage updating. Technical Report WL-TR-93-1146, Wright-Patterson Air Force Base Ohio: Wright Laboratory, Defense Technical Information Center, Cameron Station, Alexandria, VA 22304–6145, 1993.

[2] Shaul Druckmann and Dmitri B Chklovskii. Neuronal circuits underlying persistent representations despite time varying activity. Current Biology, 22(22):2095–2103, 2012.

[3] John Gibbon and Russell M Church. Representation of time. Cognition, 37(1-2):23–54, 1990.

[4] Thiago S Gouvêa, Tiago Monteiro, Asma Motiwala, Sofia Soares, Christian Machens, and Joseph J Paton. Striatal dynamics explain duration judgments. Elife, 4:e11386, 2015.

[5] Matthew T Kaufman, Mark M Churchland, Stephen I Ryu, and Krishna V Shenoy. Cortical activity in the null space: permitting preparation without movement. Nature neuro-science, 17(3):440–448, 2014.

[6] João D Semedo, Amin Zandvakili, Christian K Machens, M Yu Byron, and Adam Kohn. Cortical areas interact through a communication subspace. Neuron, 102(1):249–259, 2019.

[7] Reza Shadmehr, Helen J Huang, and Alaa A Ahmed. A representation of effort in decision-making and motor control. Current biology, 26(14):1929–1934, 2016.

[8] David Sussillo, Mark M Churchland, Matthew T Kaufman, and Krishna V Shenoy. A neural network that nds a naturalistic solution for the production of muscle activity. Nature neuroscience, 18(7):1025–1033, 2015.

[9] Richard S Sutton and Andrew G Barto. Reinforcement learning: An introduction. MIT press, 2018.

